# Targeted Chromosomal Sequencing of Wild Bonobos Identifies a Genetically Distinct Subpopulation East of the Lomami River

**DOI:** 10.64898/2026.01.08.698381

**Authors:** Mar Crego-Walters, Sebastian Cuadros-Espinoza, Irune Ruiz-Gartzia, Sojung Han, Nuria Hermosilla, Philippe Helsen, Peter Frandsen, Alexandra Maria Brumwell Prats, Pol Alentorn-Moron, Claudia Fontsere, Marina Alvarez Estape, Muzungu Ngofuna, Claude Monghiemo, Fabian Leendertz, Jo Thompson, David Fasbender, Paula Dieguez, Albert Lotana Lokasola, Colin Brand, Jean-Bosco Ndjango, Alexander V. Georgiev, Jasmin A. Giles, Weimin Liu, Yingying Li, Paul M. Sharp, Zjef Pereboom, Aida M. Andrés, Martin Kuhlwilm, Ilan Gronau, Hjalmar Kuehl, Erin G. Wessling, Victor Narat, Martin Surbeck, John A. Hart, Terese B. Hart, Christina Hvilsom, Michael Krutzen, Jeroen Stevens, Beatrice H. Hahn, Esther Lizano, Javier Prado-Martinez, Tomas Marques-Bonet

**Author notes:** Corresponding Authors: Javier Prado Martínez, Tomàs Bonet Marqués.

## Abstract

Bonobos (*Pan paniscus*), an endangered species, have for decades been genetically understudied, partly due to difficulties in obtaining high-quality samples. The study of their genome is important not only for understanding their evolution, but also for improving conservation efforts, including population management, diversity and inbreeding assessment, and tracking rescued individuals to combat illegal wildlife trafficking.

Here, we use chromosome 21 target capture data from 156 non-invasively collected faecal samples from wild bonobos to perform a comprehensive analysis of their population structure. We confirm the existence of three previously suggested subpopulations identified here as Western, Central and Eastern bonobos which are defined by natural barriers of gene flow such as the Lomami River. By estimating levels of inbreeding, diversity and differentiation, we find support for isolation of mainly Western and Eastern populations and add information on the dispersal routes of their ancestors. We infer split times and separation of these populations and apply a genetic framework to geolocalize samples of unknown origin, showing that locations of their potential origin can be estimated with a precision of up to a median of ∼50 km. Our study provides valuable insight into the evolution and population structure of bonobos and reveals how rivers act as strong barriers between populations. It also offers resources for conservation efforts and highlights the need to monitor bonobo populations more closely, in particular isolated ones.

**Impact:** Bonobos have been difficult to study genetically due to their remote forest habitat and endangered status. Here we used non-invasive sampling and chromosome 21 target capture sequencing to perform the most detailed analysis to date of their population structure. We find three main subpopulations and genetic differentiation influenced by river barriers, especially with populations found on the Eastern side of the *Lomami* river. This data will be useful for identifying the geographic origin of confiscated samples and thus aid bonobo conservation efforts.

## Background

Earth’s biodiversity is currently experiencing a mass extinction, with estimates suggesting that up to a million species are currently threatened by deforestation, climate change, pollution and other human inflicted threats (Turvey & J. 2019, UN Report 2019). Bonobos (*Pan paniscus*), are one of these endangered species (Fruth et al. 2016). This great ape, the closest extant relative to humans together with the chimpanzee, inhabits rainforests south of the Congo River in the Democratic Republic of the Congo (DRC) with their range extending from the Lualaba River in the east, to the Kasai and Sankuru Rivers in the south, and the Lake Tumba and Lake Mai-Ndombe regions in the west (Prüfer et al. 2012; Hickey et al. 2013; Mao et al. 2021). They naturally live in social groups and are mostly frugivorous; however, they are known to occasionally consume insects, fish and small mammals (Furuichi 2009). Bonobos differ in their social behaviour from chimpanzees in that females occupy high dominance ranks within society. Bonobos also exhibit reduced occurrence of intense aggression, and neighbouring groups can associate peacefully (Hohmann et al. 2003; Serckx et al. 2014; Gruber & Clay 2016; Furuichi 2020; Samuni & Surbeck 2023; Surbeck et al. 2025). However, bonobos, like chimpanzees, also have female-biased dispersal with females usually leaving their original group at an early pubertal stage (Toda et al. 2022). Female alliances dominate mating strategies and food allocation, and they are maintained by genital rubbing which reduces social tensions (Lacambra et al. 2005; Fruth et al. 2013; Reinartz et al. 2013; Gruber & Clay 2016).

While initially misidentified as chimpanzees prior to 1933, studies of this species have been hampered by their natural occurrence in remote forest regions of the DRC, a country with historical civil unrest, which in turn has accelerated poaching, illegal wildlife trade and deforestation (Coolidge J. R. 1933; IUCN 2012; Hickey et al. 2013; Ilunga Kalenga et al. 2019; Ogunnoiki 2019; Hoffmann et al. 2020; Molinario et al. 2020). Genetic studies aimed at defining the population structure of bonobos have either been limited by small high-coverage datasets from few individuals or used exclusively mitochondrial DNA (mtDNA) from faecal samples, limiting inferences to the maternal lineage (Eriksson et al. 2004; Fischer et al. 2011; Prüfer et al. 2012; Kawamoto et al. 2013; Teixeira et al. 2015; Medkour et al. 2021; Han et al. 2024).

Despite these difficulties, previous research has provided some understanding of the evolution of this species. Analysis of whole genome sequences from captive individuals determined that bonobos and chimpanzees split approximately 1.5 – 2 Mya. These studies also documented subsequent introgression events between these two species as well as the existence of an ancestral ape ghost population that has since gone extinct (De Manuel et al. 2016; Han et al. 2019; Kuhlwilm et al. 2019; Han et al. 2024). Furthermore, non-invasive sampling permitted a first analysis of bonobo population structure in the wild. Analysing faecal mtDNA sequences, two previous studies reported that the Lomami River, a major North-South river, represents a geographical barrier to bonobo gene flow. This is not apparently the case with the other West-East flowing rivers (Eriksson et al., 2004; Kawamoto et al. 2013). These studies also reported six mitochondrial haplogroups and proposed an early population split between the ancestors of bonobos now found on the two sides of the Lomami River (Takemoto et al. 2015; Takemoto et al. 2017). More recently, Han et al., confirmed population substructure using 20 high-coverage exomes, identifying three differentiated populations. Genetic differentiation, inferred divergence times and migration probabilities pointed to “Far-Western” and “Western” being more closely related to each other than to “Central” groups (Han et al. 2024).

Nevertheless, geography and the mtDNA, and differentiation at the major histocompatibility (MHC) locus *Papa-B* all suggest the presence of a third Eastern cluster that remains uncharacterised at the nuclear level (Takemoto et al. 2017; Wroblewski et al. 2023).

Genomic analyses are also important for conservation efforts because they provide insight into size, diversity and ecology of free-ranging wild populations (Foster et al. 2021; Lynggaard et al. 2022; Wei et al. 2024). They can also be used to inform inbreeding levels (Fitzpatrick et al. 2020; Dussex et al. 2021), identify the species and subspecies of certain samples (Fontsere et al. 2022), and track the illegal wildlife trade (Wasser et al. 2022). All of this can be useful information for bonobo conservation projects which impacts local ecology since this species is an important contributor (Beaune et al. 2013). Furthermore, the use of non-invasive faecal samples has enabled the obtention of DNA samples that are easier to collect and less damaging to individuals; crucial if what we aim for is data across widespread areas for the species’ conservation with help from local people (Kawamoto et al. 2013; Bowie et al. 2021; Fontsere et al. 2022). Here, we perform an extensive survey of free-ranging bonobo populations using non-invasive faecal samples across the entire DRC. We employed target capture of chromosome 21, due to its capture efficiency tested in previous research, to infer population structure and connectivity and to test whether individuals could be traced to particular geographic locations (Fontsere et al. 2022). We find a strong population substructure comprising at least three distinct groups in the west, central and eastern part of their range.

## Results

Faecal samples from wild-living bonobos were obtained from existing specimen banks as previously reported (see Supplementary table S5), or newly collected from known bonobo populations in Kokolopori in the DRC (see fig. 1). Samples were selected from 11 field sites throughout the DRC, with particular emphasis on geographic locations that have previously been underrepresented in genetic analyses. Four of these sites were located east of the Lomami River, including an established research site termed TL2, which since 2017 encompasses the Lomami National Park and buffer zone (Frankfurt Zoological Society 2013). This site was named after the rivers *Tshuapa*, *Lomami* and *Lualaba*, where samples were available from both sides of the Lomami River (Lukuru Wildlife Research Project 2009). Of a total of 272 samples, 156 passed both laboratory and bioinformatic Quality Control (QC) with an average depth of coverage of 3.46x and were suitable for analysis using chromosome 21 (see *Methods* and Supplementary figs. S1 to S7 and table S1 for further details).

**Fig. 1:**
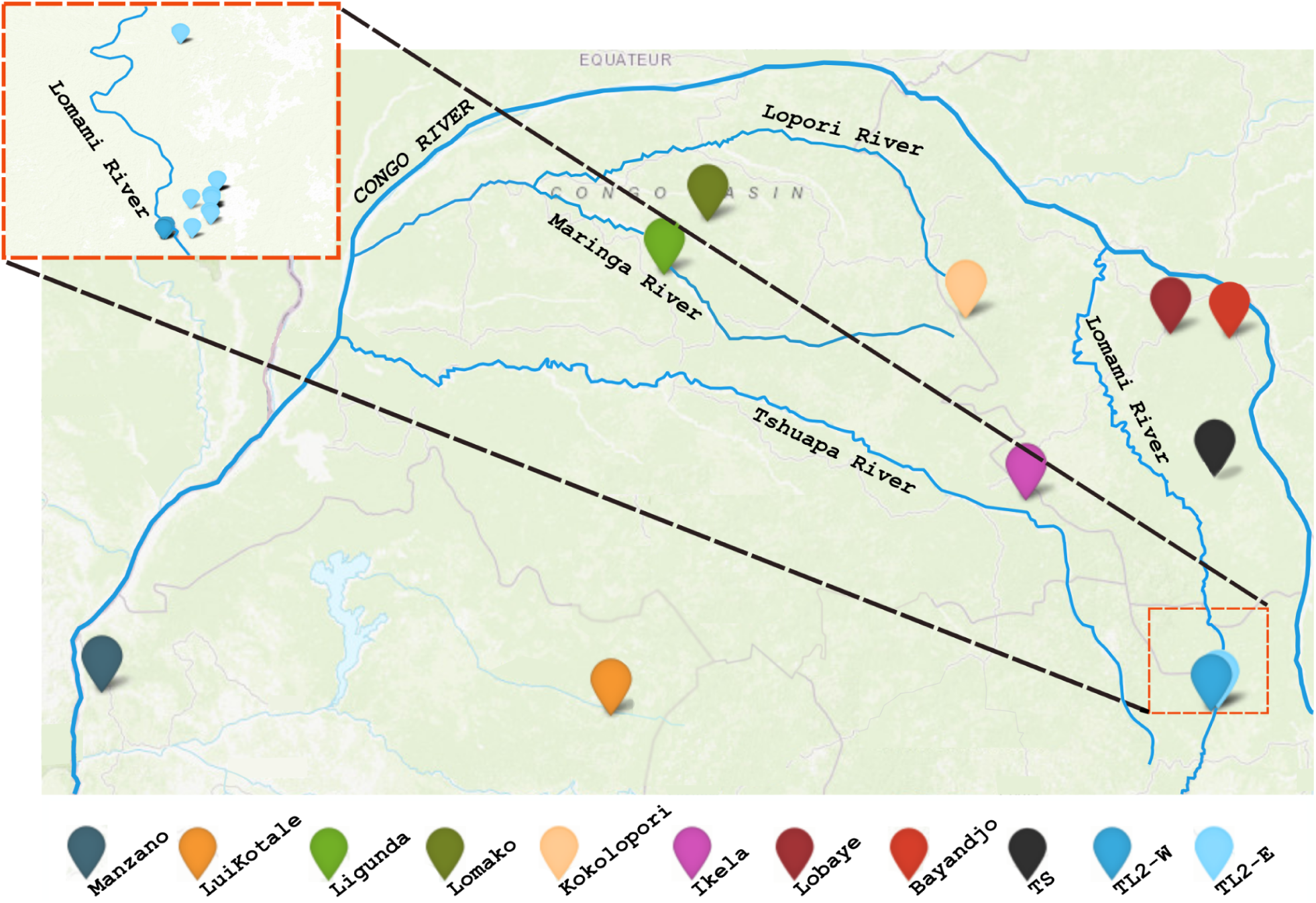
Sampling sites. The main panel shows a map in the DRC with names of sites from which samples were collected; top left square shows Lomami River separation between TL2 west and east samples. Sample size after quality filtering was: Manzano (n=11); LuiKotale (n=15); Lomako (n=18); Ligunda (n=4); Kokolopori (n=41); Ikela (n=31); TL2-W (n=6); Bayandjo (n=1); TS (n=1); Lobaye (n=1); TL2-E (n=27).

### Population Structure in Bonobos

We performed multiple Principal Component Analyses (PCA) using PCAngsd (Meisner & Albrechtsen 2018), on 156 samples with depth of coverage ≥ 0.1x. The analyses revealed that bonobo population structure generally aligns with geographical distribution forming a three-population structure which has been previously reported using mtDNA and autosomal DNA(Eriksson et al. 2004; Takemoto et al. 2017; Han et al. 2024). Specifically, Central bonobos from Kokolopori, Lomako, Ligunda, Ikela form a cluster while those from Manzano (West) and populations to the east of the River Lomami (East) drive principal components 2 and 1 respectively and form distinct clusters. Bonobos from LuiKotale cluster in between Central and West populations (see fig. 2a and Supplementary figs. S8-S9). Beyond this population structure, the PCA adds new insights, particularly regarding bonobos located between the West and East banks of the Lomami River (referred to as TL2-W and TL2-E, respectively). Despite their geographical proximity to Eastern bonobos, TL2-W individuals exhibit greater genetic similarity to Central bonobos. Furthermore, we performed an additional PCA incorporating published whole-genome bonobo data (Prado-Martinez et al., 2013) (figs. 2a & 2b). The overall population structure remains largely consistent with the addition of these samples. One exception is ‘LB502’, a bonobo born in captivity, which shows approximately 50% mixed ancestry between the Central and Eastern populations (figs. 2b & 2c). Another sample, ‘Hortense’, clusters with Eastern bonobos; however, due to limited north-eastern sample data (i.e 3 samples), it is not possible to determine whether it originates from the northeastern or southeastern TL2 region (figs. 1 & 2b). Whole-genome samples which do not clearly cluster with our site-specific samples—such as the case of ’Natalie’—may stem from their origin in nearby locations (e.g., Lac Tumba, in the west), from differences in sample type (faecal and blood), or from batch effects due to subtle differences in sequencing protocols (Tom et al. 2017). Additional PCAs using only Central bonobos with and without TL2-W also indicate a smaller scale of structure of certain Central populations, particularly in clustering Kokolopori and Lomako apart (see Supplementary figs. S10-S11).

**Fig. 2:**
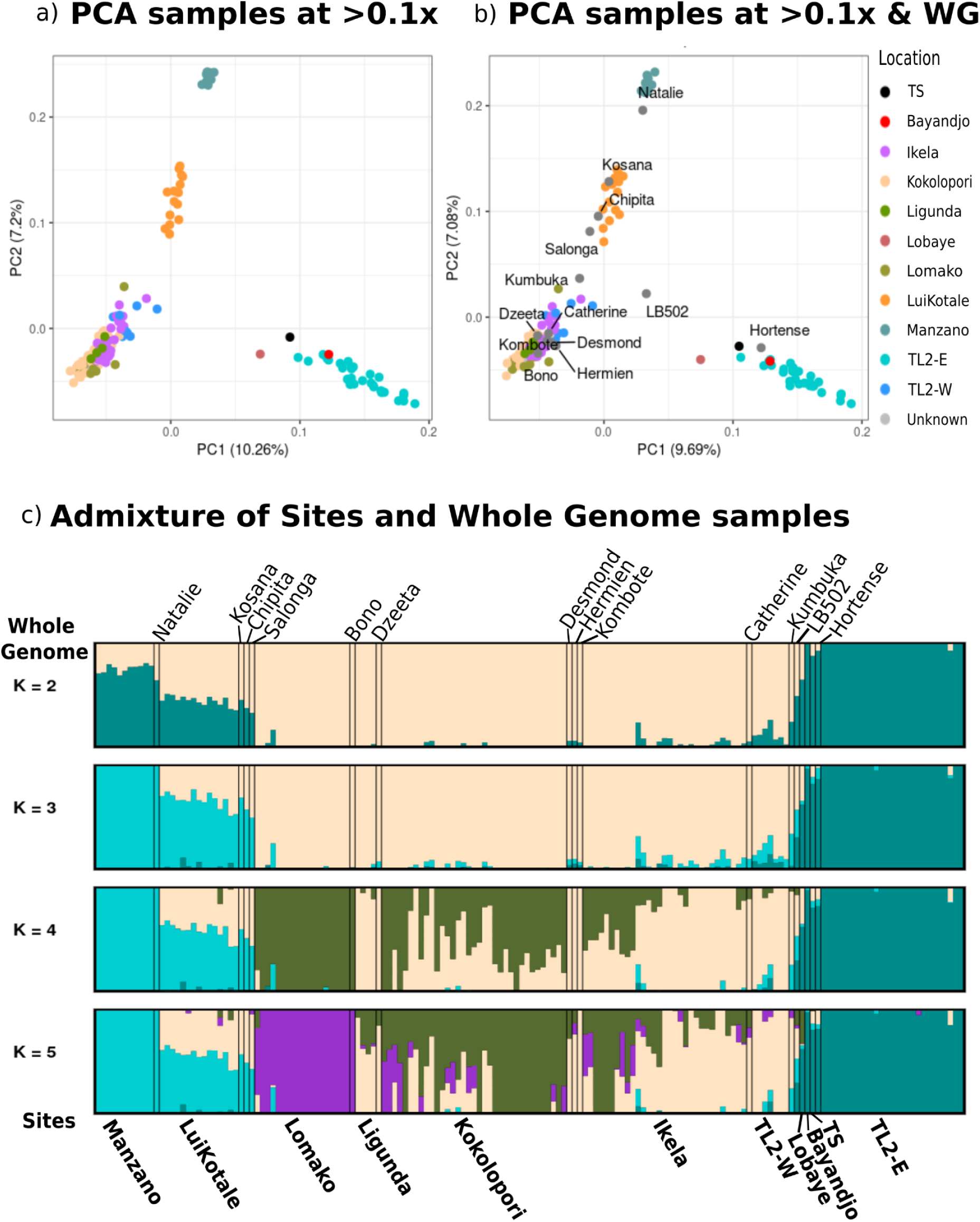
Principal Component Analysis and Admixture results using PCAngsd and NgsAdmix respectively (Meisner & Albrechtsen 2018; Skotte et al. 2013, Supplementary table S1): a) All bonobo samples which have been filtered, passed QC and with depth of coverage ≥ 0.1x; b) All bonobo samples with ≥ 0.1x and Whole Genome (WG) bonobo samples in grey from the Prado-Martinez et al. 2013 paper (n=13); c) Bar-plot of admixture values per individual grouped per site at ≥ 0.1x depth of coverage (bottom for site names) along with whole genome samples from Prado et al. 2013 (top for sample identifications).

We further explored the genetic structure of bonobos using NgsAdmix (Skotte et al. 2013; Behr et al. 2016) on 156 samples with ≥ 0.1x depth of coverage (fig. 2c and Supplementary fig. S12). Inferred K was calculated using two methods: NgsAdmix and Clumpak (see Methods; Supplementary figs. S13 & S14). The K value given was between 2 and 3 depending on method used (see Methods).

In terms of admixture we observe that Manzano, Central populations and TL2-E show their own ancestry components (see fig. 2c). On a more detailed level, Manzano individuals share genetic ancestry with LuiKotale, the closest site with no natural barriers (see fig. 1 & 2c). Central bonobos from Lomako, Ligunda, Kokolopori and Ikela share genetic ancestry, coinciding with results from the PCA. The TL2-W site also shares substantial ancestry with those central populations. Lomako has its own ancestral component at K=4 (fig. 2c & Supplementary figs. S10-S11).

We next assessed each population’s genetic diversity, heterozygosity, inbreeding and relatedness. We calculated levels of heterozygosity using ANGSD (version 0.935, Korneliussen et al. 2014) and found that Central bonobos exhibit higher levels of heterozygosity, whereas non-Central bonobos such as TL2-E and Manzano are lower (fig. 3a). We should note that after filtering for 5x depth of coverage, Ikela only has one sample with a depth of coverage of 4.94x which was included for informative purposes but could imply bias in this analysis for this site (see table S1 in Supplementary).

**Fig. 3:**
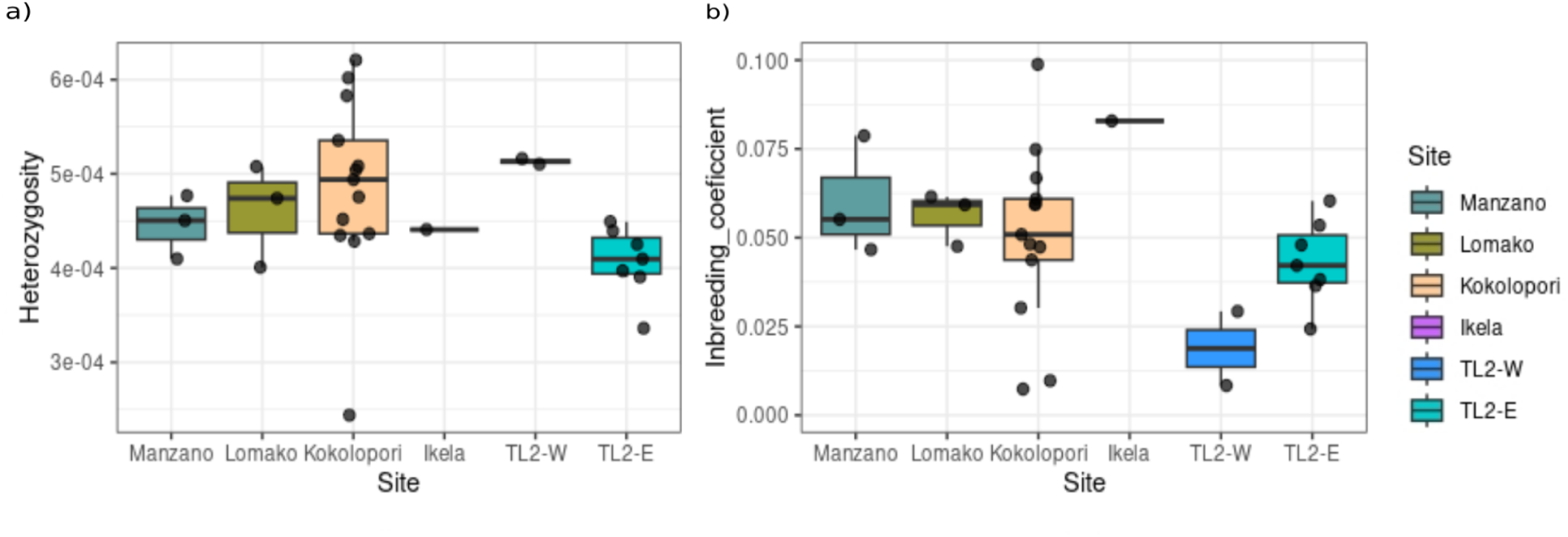
Heterozygosity and inbreeding evaluation (see Supplementary table S1 for number of samples): a) box-plot of heterozygosity levels per site at 5x (n=29) depth of coverage or higher; b) box-plot of inbreeding coefficient levels per site at ≥5x (n=29) depth of coverage.

Additionally, by comparing heterozygosity levels with different bins of depth of coverage in Supplementary fig. S15, we conclude that we have limited resolution to assess heterozygosity in these non-invasive samples, due to low depth of coverage which can greatly influence results (Fontsere et al. 2022). We thus focus on general comparisons between populations rather than true values for the different samples and locations.

For inbreeding, we used Fhat3 estimate from PLINK (version 1.9, Chang et al. 2015, see Methods) and found elevated inbreeding coefficients in Ikela,Manzano and to a lesser level in Lomako, but lower levels in TL2-W, close to 0 (see figure 3b). Bonobos have been previously seen to have relatively high levels of inbreeding in the Western populations so in the case of Manzano, our results match previous results (Han et al. 2024). Although the variation in Kokolopori is noticeable, we were unable to compare it to the rest of sites due to the low number of high quality samples in these, which in turn offers low statistical power for a formal analysis of variance.

Inbreeding values for Ikela, despite being compatible with the heterozygosity values above, should be taken with caution as it has been evaluated as an individual sample (see fig. 3b & Supplementary table S1). We also analysed related individuals using NgsRelate v.2 (Korneliussen & Moltke 2015) and found a total of nine first-degree relatives although relatedness per site cannot be directly interpreted due to sampling bias (see Supplementary fig. S16 to S18). The related individuals were removed prior to the aforementioned analysis to avoid bias (Fontsere et al. 2022).

### Population differentiation and divergence

#### *F_ST_* population values

In order to analyse differentiation between populations, we calculated *Hudson’s F_ST_* using ANGSD at ≥0.9x depth of coverage and Plink 1.9 at 5x depth of coverage which show agreement within the bonobo trends (fig. 4 and Supplementary fig. S19 respectively; see for sample size in this case, *Methods*). We compared *F_ST_* values between bonobo sites, and bonobo areas with chimpanzee subspecies data published in Prado-Martinez et al. 2013 and De Manuel et al. 2016 (see figures 4 & Supplementary fig. S19).

**Fig. 4:**
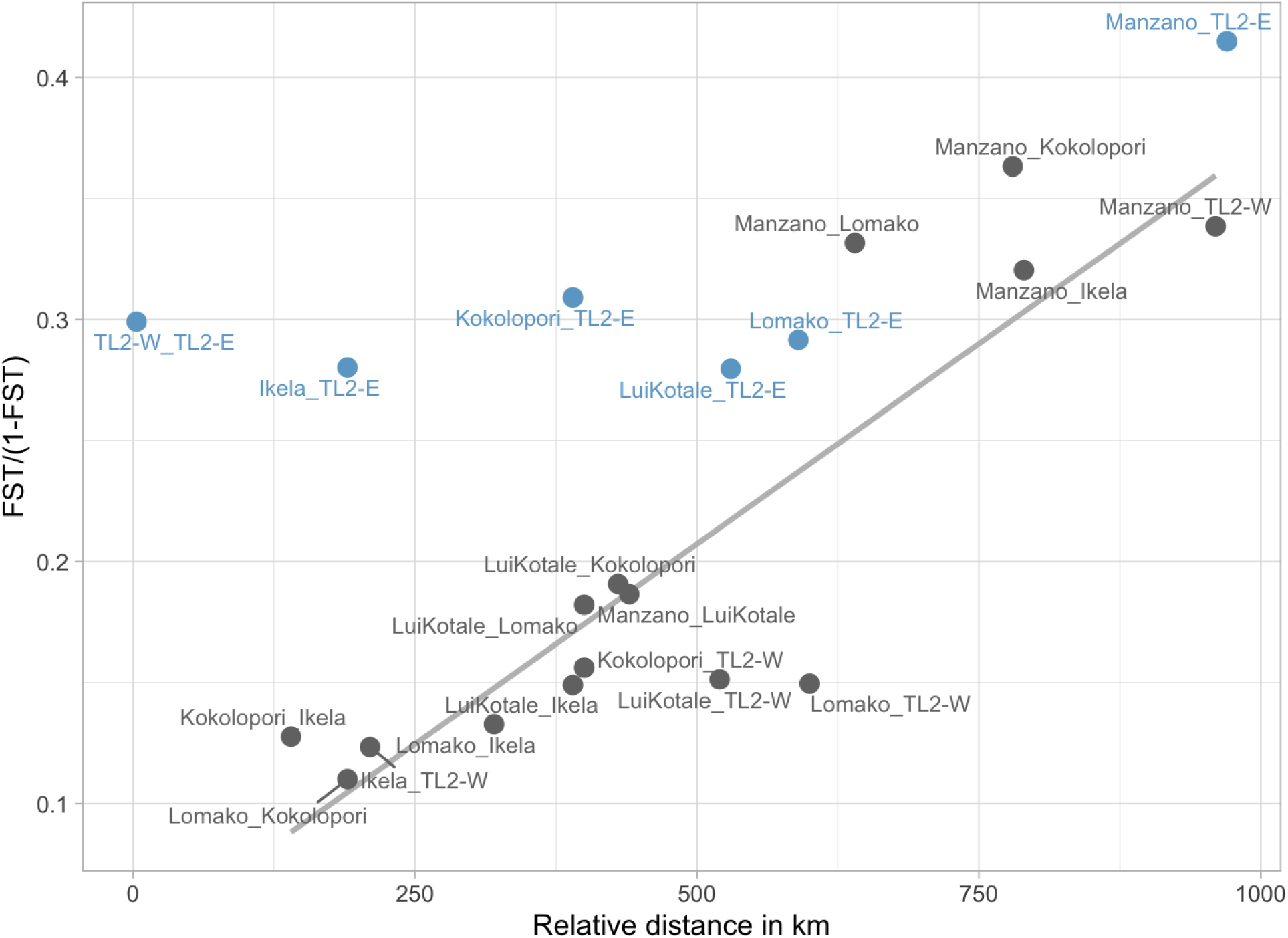
Scatterplot of *F_STst_/(1-F_ST_)* sites index versus distance between sites in km. The correlation line is for all points except those including TL2-E (in blue). Genetic differentiation on x axis and distance in km on y axis.

Sites with highest pairwise differentiation are Manzano and TL2-E (see below fig. 4). These two are the most isolated and geographically distant sites in the westernmost and easternmost areas respectively (see fig. 1). Although Manzano has higher differentiation with most sites, it presents low differentiation with the closest site: LuiKotale. Pearson correlation values between *F_ST_/(1-F_ST_)* and distance for all sites including TL2-E was of R² = 0.619 (with *p-value* = 0.0027). When we remove TL2-E, this value rises to R² = 0.87 (with *p-value* = 1.828e-05).

Therefore, TL2-E is the only site that does not follow isolation by distance and can be seen as an outlier from the rest of sites, especially with the closest ones in terms of distance: TL2-W and Ikela. Interestingly, pairwise comparisons between either TL2-E or Manzano and a Central population yield very similar values of *F_ST_* regardless of geographical distance, which in turn could indicate a basal separation for both Manzano and TL2-E from Central Bonobo populations.

When comparing *F_ST_* values of chimpanzee subspecies and bonobo areas, we observe a similar correlation between genetic differentiation and geographical distance, although this is overall less pronounced in bonobos (Supplementary fig. S19 & table S2; Fontsere et al. 2022). Distance between the two furthermost chimpanzee subspecies in terms of kilometres, *Pan troglodytes verus* and *Pan troglodytes schweinfurthii*, is of ∼2000km (Lester et al. 2021; Takoukam Kamla et al. 2021). For bonobos, the distance between Western and Eastern bonobos is ∼750km (Supplementary table S3; Hickey et al. 2013). *F_ST_* values for bonobo areas indicate higher genetic differentiation between the furthest located bonobos, i.e. Western and Eastern, but also between Western and both Central bonobos (see Methods for sample size and Supplementary fig. S19, right). In the case of Western bonobos, it confirms the already existing theory that Western bonobos are genetically isolated and follow the isolation by distance trend (Han et al. 2024). This showcases these effects which are already apparent between chimpanzee subspecies and greatly affects differentiation.

#### Demographic modelling using G-PhoCS

Divergence time between the West and Central bonobos was previously estimated at ∼145 kya (Han et al. 2024). However, the phylogenetic relationship between the three bonobo populations (Eastern, Central, and Western) has been unclear so far. A tree based on mtDNA haplogroups suggested that the Eastern bonobos branched out from the common ancestor of the Western and Central bonobos (Takemoto et al. 2017; Han et al. 2024). Nevertheless, a neighbour-joining tree inferred from whole-genomes in another study did not suggest a significant structure within the bonobo populations (de Manuel et al. 2016). In order to estimate split times and migration rates between the bonobo populations, we used G-PhoCS, a Bayesian inference method that utilizes a coalescent sampling of local genealogies to infer population divergence times, migration rates, and effective population sizes (Gronau et al. 2011; Han et al. 2024). This method is designed to produce reliable estimates from few (or only one) high-quality genomes per population. Based on the results above, we selected the Western, Central and Eastern populations as the main lineages to analyse and exploit whole-genome data (Prado-Martinez et al. 2013). The geographic origins of individuals contributing these genomes were previously inferred from their mtDNA (Han et al. 2024), as well as their clustering in the PCA and ADMIXTURE (figure 2). As there was only one individual whole-genome available representing the Eastern population (Hortense), one individual per population was selected. A western chimpanzee was used as an outgroup, as most likely no recent gene flow between bonobos and this subspecies occurred (De Manuel et al. 2016; Brand et al. 2022).

In order to resolve the order of divergence from the common ancestor, we used three-population models, each including two of the three bonobo populations and chimpanzee as an outgroup (Supplementary figure S22). We first considered models without migration, with results suggesting similar divergence times for the three pairs of bonobo populations (Eastern-Central, Eastern-Western, Central-Western) at around 238 thousand years ago (fig. 5, Supplementary table S4). When gene flow between populations is allowed after their split, the divergence times increase, but they remain similar across the three pairs, at 493 kya with very high migration rates of up to 20% (Supplementary table S4). We note that across all models with and without gene flow, the divergence time between bonobos and chimpanzees was estimated consistently to be ∼1.4 Mya (1.37-1.43 Mya), which is within the range of previously inferred estimates (Prüfer et al. 2012; De Manuel et al. 2016; Kuhlwilm et al. 2019; Han et al 2024). We conclude that at this stage we cannot disentangle the order of splits between these bonobo groups. This is likely due to a short time span during which divergence occurred, probably not as a sharp divergence event, and possibly followed by high levels of post-divergence gene flow. However, these factors cannot be determined with the data available.

**Fig. 5:**
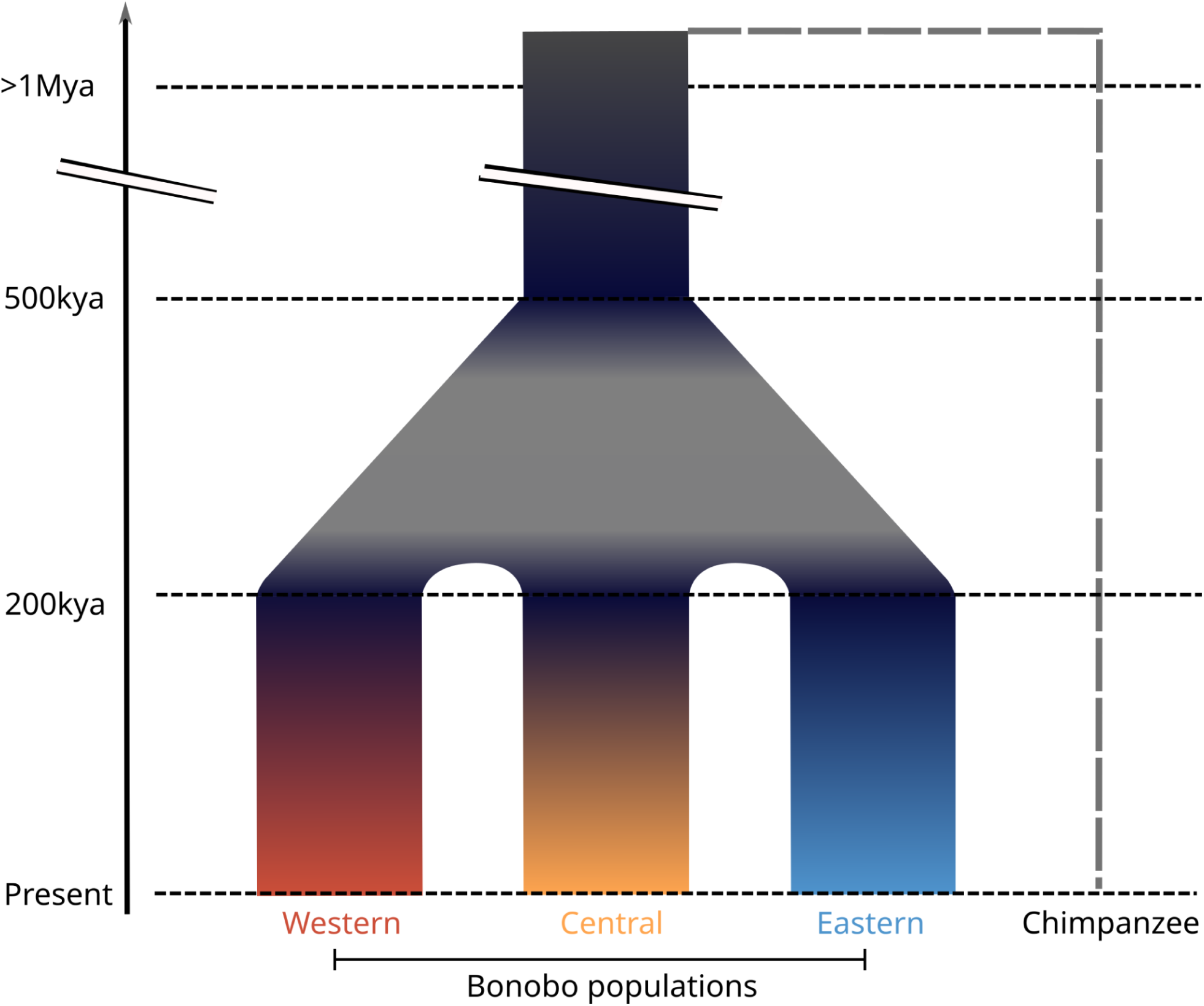
Conceptual model of bonobo population divergence, based on G-PhoCS results. Split times shown on the y axis, population labels on the x axis. A chimpanzee mutation rate (μ=0.64e-9, Langergraber et al. 2012; Besenbacher et al. 2019) was used for the calibration. Further details in fig. S22 and table S4.

Our analyses suggest a lower bound of 238 kya for the divergence of the three bonobo populations, which is deeper than previous estimates between Western and Central bonobos (∼145kya, Han et al. 2024). Similarly, the bonobo-chimpanzee split time of 1.4 Mya is deeper than the previous estimate of 1.2 Mya. These differences likely result from using a different reference genome and improved filtering procedures (see *Methods*), despite using the chimpanzee mutation rate (μ=0.64e-9, Langergraber et al. 2012; Besenbacher et al. 2019), which is faster than the human mutation rate (μ=0.43e-9) used in Han et al. 2024 for calibrating G-PhoCS estimates.

#### Inference of geographic origins

We used the deep learning method Locator to assess whether we are able to predict geographic origin of other samples based on our chr21 sample coordinates (Battey et al. 2020). We initially tested and trained the method by using known location samples from our high coverage chr21 (n=29) dataset (see fig. 6 and Supplementary excel *Locator_results.xlsx*) and thereafter predicted the origin of the whole genome samples from Prado et al. 2013 (Supplementary fig. S21). All analyses were performed with replicate models on bootstrap samples and iterating over validation samples, i.e. samples with coordinates, as recommended by developers (Battey et al. 2020).

**Fig. 6:**
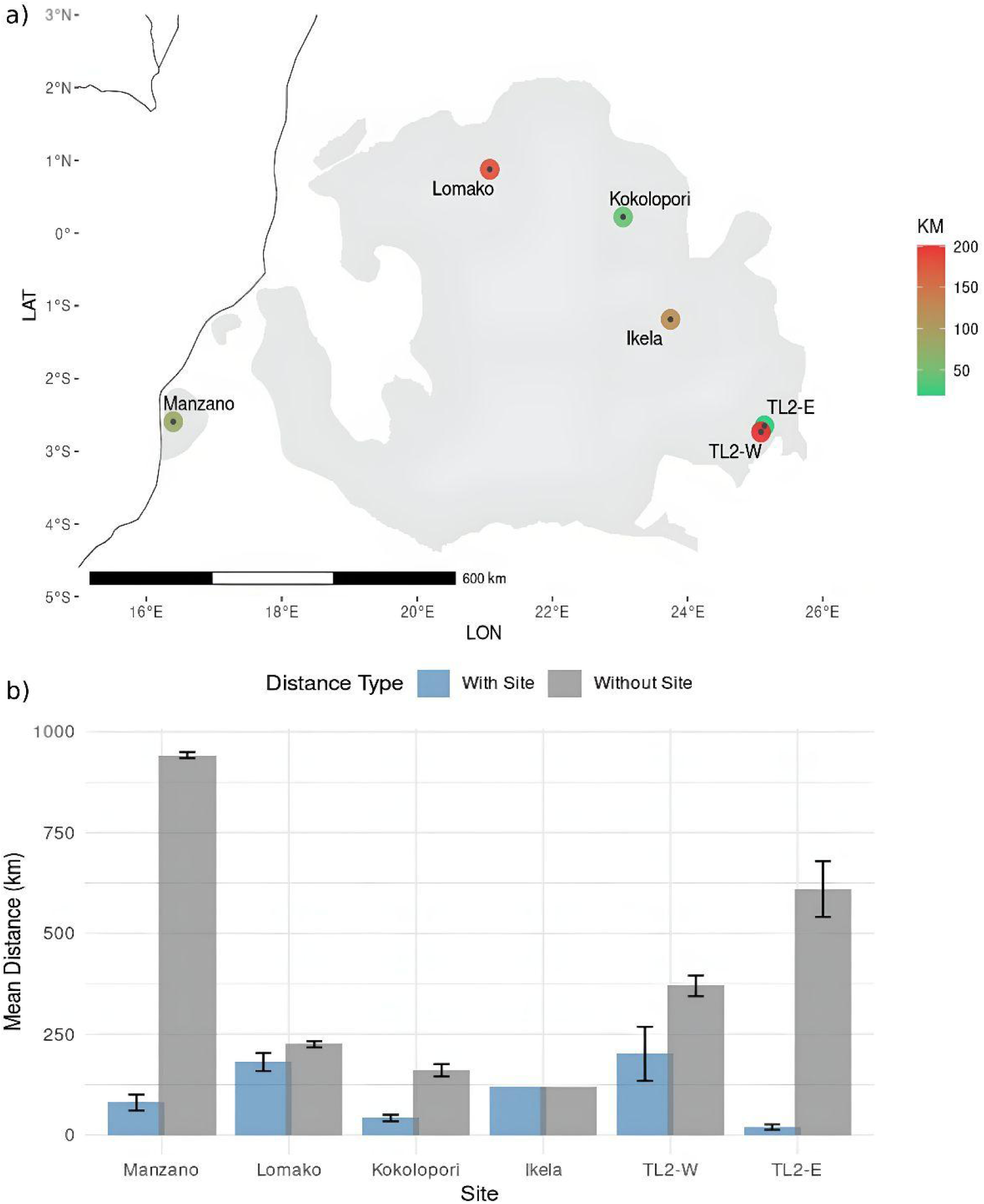
Results from geolocalization using our chr21 high coverage dataset (see Supplementary table S1 for number of samples): a) Map with coloured error rate of average km from real site in Locator predictions when removing single samples; grey area is the bonobo habitat range; b) Barplot of kilometres’ distance from real coordinates of all our tests ***with site***, meaning with other same-site samples (blue) included in the analysis and ***without site***, meaning without any more site samples (grey) included (i.e. the other site samples were otherwise removed). For Ikela we only have one sample at 5x, thus once removed there are no more site samples.

For the unknown whole genome samples, we removed hybrid sample LB502, and samples that had low depth of coverage and only single end data. We were then able to run Locator for 9 of these samples and estimate the collection location based on these results and those from the previous PCA and Admixture analysis (see Supplementary fig. S21).

We were able to approximate location by a median of 53 km and a maximum average of 268 km and a minimum average of 5.06 km distance from real locations in the initial tests which, considering the 5km range of some bonobo groups, is an important approximation (Supplementary excel *Locator_results.xlsx;* Eriksson et al. 2004). To test performances further, we ran Locator without other samples from the same site and found that in these cases, it approximates the location to the next nearest site with known coordinates and in the case of TL2-E and Manzano, error rates are much higher when we have no more same-site samples (see fig. 6). The sites Lomako and TL2-W also showed higher error rates, likely due to their low sample sizes and lower population differentiation with other Central populations. In particular, TL2-W also may be confounded by the small geographic distance to TL2-E which, given the strong differentiation between these populations, might distort the accuracy of this prediction. These tests show the need to sample more populations for increasing predictions and accuracy, especially those found in the peripheral populations of bonobos.

## Discussion

We utilized 156 out of an initial 272 non-invasively collected faecal samples for our analysis (see Supplementary file: *Metrics_final.xlsx*). This highlights a critical consideration when working with non-invasive samples, as many fail quality control (QC) or may represent identical individuals (see Methods section; Han et al. 2025). As such, we emphasize the importance of collecting a sufficient number of samples to improve overall dataset quality, particularly in understudied regions, in our case Salonga, Ikela and TL2-W.

Despite the challenges associated with analysing these samples, our findings corroborate the existing model of three main genetic clusters in bonobos and further underscore the role of rivers in restricting bonobo gene flow through the use of chromosome 21 data (Eriksson et al. 2004; Takemoto et al. 2017; Han et al. 2024). Specifically, we demonstrate that the Western population from Manzano is highly differentiated, consistent with a previous exome study (Han et al. 2024), and establish that a population from the east of the Lomami River (TL2-E) is similarly distinct, consistent with mtDNA data (Takemoto et al. 2017). For Western populations, it is important to highlight that the Manzano area is a highly fragmented bonobo habitat which could potentially restrict gene-flow (Narat 2011; Pennec et al. 2020). Interestingly, TL2-W populations share substantial genetic variation with Central bonobos but are highly differentiated from Eastern/TL2-E populations. This indicates that the Lomami River system has functioned as a significant genetic barrier for possibly all Eastern Lomami river populations. In the case of TL2-W, it can lead to biased results when inferring origin if only few analyses such as PCA are used, i.e. it clusters with central populations such as Ikela and Kokolopori (Elhaik 2022).

Nevertheless, while the Lomami is apparently a barrier to bonobo gene flow, previous research has found that the *Plasmodium* populations at TL2-W and TL2-E are not divergent meaning it does not prevent mosquitoes from transmitting malaria parasites between the two sites (Liu et al. 2017). The overall genomic differentiation seen here between TL2-W and TL2-E bonobos reinforces the conclusion that similarity in the types of MHC alleles seen at the two sites likely reflects independent adaptation to malaria (Wroblewski et al. 2023). This suggests that, although TL2-W is genetically closer to Central bonobos, it remains sufficiently isolated from Central bonobos to exhibit similar infection patterns to TL2-E. Furthermore, our analysis could not resolve the order of divergence between the three main populations, which suggests divergence close in time or potentially even a trifurcation. The divergence of bonobos from their ancestral population into three distinct populations might have occurred due to a decrease in the Congo river discharge, allowing a series of dispersal corridors for these populations, as previously proposed (Takemoto et al. 2015). As such our new analysis adds information to the bonobo dispersal theory using nuclear DNA, although certainly not in a conclusive manner with the current data from whole genomes. If more individuals representative of these populations become available in the future, a more fine-grained demographic modeling will be an important next step, e.g. using SFS-based or other methods (Gutenkunst et al. 2009; Excoffier et al. 2021; Soni & Jensen 2025).

Understanding these river-driven population structures is crucial for conservation efforts. It not only impacts geolocalization in conservation projects—where even geographically close samples may be difficult to assign accurately due to natural barriers (see fig. 4)—but also influences levels of heterozygosity and inbreeding in populations such as Manzano and TL2-E bonobos. To improve the accuracy of sample origin identification, we applied a deep learning approach, which successfully predicted all sample locations within a median radius of 53 km from their actual origin (see fig. 6). Additionally, we would like to highlight that the strongly differentiated populations of Manzano and TL2-E warrant particular attention, as their genetic isolation may heighten their vulnerability to environmental pressures (Theissinger et al. 2023; Han et al. 2024). The TL2-E population in particular exhibits significant genetic differentiation from other groups due to the Lomami River which limits connectivity with Central and Western populations and, most likely, gene-flow.

## Conclusions

This study adds an extensive genomic dataset from non-invasively collected samples, focusing on nuclear DNA. Our results highlight the genetic differentiation of bonobo populations, especially in the case of Eastern and Western populations and adds information to the question of ancestral origin of Bonobo populations, hinting at a more complex dispersal theory. In addition, our analysis demonstrates the possibility of geo-localization of unknown samples which can inform future conservation projects by identifying the most likely geographic coordinates based on genetic variation. However, we stress the need for additional samples from other regions, particularly South-Western (Salonga area and south of the Lukenie river) and Western (Lac Tumba) bonobos, to refine these predictions further (see *Inference of geographic origin* section; Hickey et al. 2013; Pilbrow & Groves 2013). Future research should focus on male lineage samples to further understand ancestor dispersal, finding informative variants, a formal analysis of variance of heterozygosity and inbreeding values between sites, identifying markers to differentiate populations within the Central bonobo group, alongside leveraging advancements in AI and portable sequencing technologies to speed up data collection, lower costs, and expand the availability of research methodologies in local settings (Watsa et al. 2020; van Oosterhout 2024).

Finally, we emphasize the importance of integrating conservation genomics with rehabilitation initiatives for wild bonobos to ensure the survival of this endangered species. Our findings provide valuable insights into relative genetic diversity across populations, identify potential individuals with admixed components, and enable the non-invasive determination of wild individuals’ origins. This could even lead to pinpointing such origins to the specific side of a river with approximate coordinates. This approach not only supports conservation efforts but also minimizes harm to individuals, aligning with ethical research practices.

## Methods

### Sample selection and laboratory methods

A total of 272 wild Bonobo samples were collected from different locations and providers (see Supplementary *Metrics_final.xlsx*; Supplementary Table S5). From these samples we performed an initial shallow shotgun sequencing following the BEST protocol as explained in the Carøe et al. 2018 paper to determine the amount of host DNA (hDNA). Prior to this, DNA samples were sonicated to obtain fragments of approximately 200 base pairs using the Q800R3 sonicator (Qsonica, USA). The sonication was performed at 40% amplitude in pulse mode (15 seconds on / 15 seconds off) for a total active sonication time of 15 minutes. Three samples with less than 0.01% hDNA were discarded whilst the rest were sequenced using a previously developed target capture method of chromosome 21 (see table 1 and *Metrics_final.xlsx* in Supplementary; Fontsere et al. 2021). Chromosome 21 had been used in previous studies due to its size which increases target capture efficiency in faecal samples and its relatively high population genomic information content. Samples were pooled according to hDNA content (Hernandez et al. 2018), and divided into aliquots to then capture chromosome 21 as performed in Fontsere et al 2021, for the subsequent hybridizations. Captured libraries were sequenced on a NovaSeq 6000 system with 2 x 150 paired-end reads. We also excluded samples that contained more than 1% human contamination (see Supplementary table S1).

### Data processing and filtering

We processed the data to demultiplex libraries belonging to the same hybridization pool using Sabre (https://github.com/najoshi/sabre) and checked the quality of the fastq files using FASTQC (Andrews 2010). Reads were then trimmed to a minimum of 30bp with fastp version 0.23.2 (Chen et al. 2018).

Paired-end reads were then aligned to the human genome Hg38 (GRCh38, Dec.2013 (GCF_000001405.26)), with a minimum length of 35bp using BWA (version 0.7.17). Duplicates were removed using PicardTools (version 2.26.10) (http://broadinstitute.github.io/picard/) and additional filtering of the reads was done using SAMtools (version 1.15). We then intersected the reads of Chr21 using BEDTools version 2.30.0. In addition, we performed a human contamination test using HuConTest (Kuhlwilm et al. 2021) and discarded samples that had more than 1% human contamination (see Supplementary figures S1 to S4 & excel *Metrics_final.xlsx*).

Average depth of coverage of the target space was calculated as the number of bases in the target region divided by the size of chromosome 21 (see Supplementary excel *Metrics_final.xlsx*). We then excluded samples that had a lower depth of coverage than 0.1x, leaving 156 samples with an average depth of coverage of 3.46x (see Supplementary table S1).

We compared depth of coverage per site with hDNA per site and calculated library complexity, capture specificity and enrichment factor for each pool as an additional QC measure (see Supplementary figs. S1 to S7 and excel *Metrics_final.xlsx*). We obtained genotype likelihoods and genotype calls using ANGSD version 0.935 (Korneliussen et al. 2014) and commands angsd -b samples.list -ref ${REF} -uniqueOnly 1 -remove_bads 1 -only_proper_pairs 1 -trim 0 -C 50 -baq 1 -minInd 10 -skipTriallelic 1 -GL 2 -minMapQ 30 -nThreads 24 -doGlf 3 -doMajorMinor 1 -doMaf 1 -minMaf 0.004 -SNP_pval 1e-6 -r chr21: -out outfile.

To determine relatedness and remove duplicated or identical twins’ samples, we used NgsRelate v.2 (Korneliussen & Moltke 2015) at a ≥ 0.3x depth of coverage (fig. 3; Supplementary fig. S16 to S18). For most analyses we filtered samples by depth of coverage of ≥ 0.1x but for those that required more high-quality data we filtered by ≥ 0.3x or ≥ 0.5x or ≥ 5x (see Supplementary table S1).

### Data analysis

Principal Component Analysis (PCA) was performed using PCAngsd (Meisner & Albrechtsen 2018), filtering samples at 0.1x (n=156) and removing duplicated or related samples (fig. 2; Fontsere et al. 2022). We performed the covariance matrix calculation using the following command: python pcangsd -b BAM.beagle.gz -e 2 --threads 64 -o BAM_chr21_covmatrix. We performed several PCAs: one with all captured samples; another, with whole genome samples from the Prado-Martinez et al. 2013 paper; and several PCAs with only Central bonobo samples with and without TL2-W (see fig. 2 and Supplementary figures S8 to S11).

To evaluate population structure, we used NgsAdmix with -minMaf 0.05 command across 50 simulations (K values up to 11) and plotted with pong at higher or equal than 0.1x depth of coverage (n=156). Inferred K was calculated by using two methods: choosing the value of K with the highest normalized mean likelihood (across runs) and the Clumpak method which chooses the value that maximizes the normalized second-order change in log-likelihood across iterations. Both methods were calculated manually in R v. 4.4.x and in CLUMPAK for comparison (see Supplementary figs. S16, S17 & S18; Skotte et al. 2013; Kopelman et al. 2015; Behr et al. 2016; Posit Team 2022; R Core Team software Version 4.4.x. 2025). When the inferred K is calculated from the normalized mean from all the Ks, the given value is 2. However, if the inferred K is calculated from the normalized median of differences from likelihood of all Ks, then the given value in both methods is 3 (see Supplementary figs. S16 & S17). We analysed samples by site and added the whole genome samples later on (see fig. 2).

To determine levels of heterozygosity, we used ANGSD (version 0.935; Korneliussen et al. 2014) with the *realSFS* command to estimate site frequency spectra. In order to avoid introducing biases due to heterogeneous coverage (Fontsere et al. 2022) and compare our results of inbreeding, we tested the results using SAMtools (version 1.15) by filtering out samples below 5x depth of coverage and downsampled the remaining ones to 5x (n=29) depth of coverage (see fig. 3a and 3b; Supplementary table S1). Command used was: angsd -i ${BAM} -anc ${REF} -ref ${REF} -uniqueOnly 1 -remove_bads 1 -only_proper_pairs 1 -trim 0 -C 50 -baq 1 -minMapQ 20 -minQ 20 -setMaxDepth 200 -doCounts 1 -GL 1 -doSaf 1 -r chr21: -out ${ID}_heterozyg | realSFS ${ID}_heterozyg.saf.idx >${ID}_est.ml (Nielsen et al. 2012).

We performed variant calling using GATK v.4.2.5. for bams with depth of coverage above or equal to 5x (n=29) of our bonobo samples and for the chimpanzee area *F_ST_* (Supplementary fig. S19), we added the published chimpanzee bams from the Prado-Martinez et al. 2013 and De Manuel et al. 2016 papers. We then combined the vcfs using CombineGVFCs and then genotyped GenotypeGVCFs command and removed sites with missing data with VCFtools v.0.1.16 using --max-missing 0.5 leaving a total of 660992 SNPs.

For inferring inbreeding, we used the same initial filtering process of ≥ 5x depth of coverage and downsampling to 5x depth of coverage as with the previous heterozygosity analysis (see section above), using the Fhat3 estimate with PLINK (version 1.9, Chang et al. 2015). We pruned the vcf containing all samples with 5x (n=29) depth of coverage (see fig. 3). Command used was: plink --vcf ${vcf} --read-freq Bonobos_downsampled.frq --set-missing-var-ids @:# --het –ibc --out Bonobos_5x.

When calculating pairwise *F_ST_* estimates between areas (see Supplementary fig. S19) and sampling sites (see fig. 4), we used PLINK (version 2, Chang et al. 2015) for a vcf with samples at 5x with --fst CATPHENO method=hudson --set-missing-var-ids @:# command options and at ≥0.9x depth of coverage ANGSD (version 0.935, Korneliussen et al. 2014) with -anc ${REF} -dosaf 1 -gl 1 and realSFS pop1.saf.idx pop2.saf.idx -P24 > pop1.pop2.ml | realSFS fst index | realSFS fst stats pop1.pop2.fst.idx command options for sample bams (Nielsen et al. 2012). Previously, we also downsampled to 4 samples per site and 11 per area, and plotted the unweighted and weighted values to a heatmap using R (Posit team 2022; R Core Team software Version 4.4.x. 2025). The option of hudson estimates *F^stHudson→(Fst^1^+ Fst^2^)/2* was chosen since this estimate is not influenced by sample sizes (see equation below from Hudson et al. 1992; Weir & Hill 2002; Bhatia et al. 2013). We then divided *F_ST_* by (1-*F_ST_*), since it is not bounded between 0 and 1, yielding a result that is better related to the geographic distance (Rousset 1997).

To infer the demographic history of the three bonobo populations, we used the Generalised Phylogenetic Coalescent Sampler (G-PhoCS), a Bayesian inference method that uses Markov Chain Monte Carlo (MCMC) sampling of local genealogies at short, putatively neutral loci in approximate linkage equilibrium to guide posterior sampling of demographic parameters (Gronau et al. 2011). G-PhoCS infers demographic parameters, such as divergence times, effective population sizes and migration rates, given a pre-specified population phylogeny. G-PhoCS can produce reliable parameters using as few as one whole genome per population. Here we used bonobo whole genomes published in Prado-Martinez et al. selected based on their PCA clustering with our chr21 samples and the admixture analysis (Population Structure in Bonobos, fig. 2b & 2c). Data was mapped to the human reference genome hg38, with joint genotype calling across the *Pan* clade, and subject to quality assessment as described in Han et al. 2025. We filtered and prepared the genotypes following the manual (http://compgen.cshl.edu/GPhoCS/GPhoCS_Manual.pdf), using default parameters. Eight quality filters were downloaded from the UCSC genome database (http://hgdownload.cse.ucsc.edu/goldenpath/hg38/database/, the last date of access: 29.01.2025), to remove known genic regions (refGene, knownGene), simple and complex repeat regions (simpleRepeat, genomicSuperDups), CpG islands (cpgIslandExt), repeat masker (rmsk), conserved regions across 30 primate species (phastConsElements30way), and to keep the loci within the synteny net between the assemblies hg38 and PanPan3 (netPanPan3). Removing conserved regions should remove most functional sites, which will likely account for effects such as background selection (Johri et al. 2021). A set of loci of 1000 bp after applying the filters was obtained, which were at least 20 kb apart from each other and had less than 20% missing data across all the individuals used in each analysis. The number of such blocks across the genome was 22,830. We then calculated mean values as point estimates and 95% Bayesian credible intervals as plausible ranges. To convert Tau and Θ values to calendar year and Ne, we used a chimpanzee mutation rate of 0.64e-9 per site per year and a generation time of 24 years (Langergraber et al. 2012; Besenbacher et al. 2019). In our case, migration rates were incorporated into the model to account for the impact of gene flow on divergence time estimates, rather than to obtain precise estimates of the migration rates themselves—an aspect that remains one of the major unresolved challenges in population genetic modeling. A migration proportion, which ranges between 0 and 100%, was calculated for each migration band by multiplying its inferred migration rate with the associated tau estimate. These proportions approximate the probability that a lineage from the target population of a migration band experienced migration. In each analysis, we considered three populations, where a western chimpanzee was used as an outgroup of two bonobo populations. We considered all three possible pairs of bonobo populations. For each pair of bonobo populations, we ran a G-PhoCS analysis without any migration band and a separate analysis with four migration bands: two between the bonobo populations and two between the ancestral western chimpanzee population and the population ancestral to the two bonobo populations. For each of the six models, we ran 10 replicate runs, which had different starting states. After discarding the first 500,000 MCMC iteration as a burn-in, the last 500 thousand MCMC interactions were used.

We used the deep learning method Locator to predict the unknown origin of the bonobo samples from Prado-Martinez et al. 2013 (Battey et al. 2020). We removed sites with missing data with VCFtools v.0.1.16 using --max-missing 1 leaving a total of 8405 SNPs as per user instructions. We trained replicate models with our 5x chr21 dataset on bootstrap samples of the vcf with iterations and looped over seeds to generate the predictions with commands --seed $i --bootstrap --nboots 50 (Battey et al. 2020). We then tested the method by picking random samples from our dataset to predict location and also in a leave-one-out validation method, which gave approximations to the real coordinates by 5.06 km difference from real location (see fig. 6, Supplementary fig. S20 & excel *Locator_test_results_final.xlsx*). We also tested by eliminating samples from specific sites and running the method to predict these locations (see Supplementary *Locator_test_results_final.xlsx*). Finally, we used nine whole genome samples from Prado-Martinez et al. 2013 and predicted their coordinates (see Supplementary fig. S21). To better represent our results we modified Locator’s output plot rscript using R (Battey et al. 2020; Posit team 2022; R Core Team software Version 4.4.x. 2025).

## Declarations

### Ethics approval and consent to participate

Not applicable.

## Consent for publication

Not applicable.

## Availability of data and materials

The datasets generated and/or analysed during the current study are available from the ENA repository with accession number PRJEB96168 [https://www.ebi.ac.uk/ena/browser/view/PRJEB96168]

## Competing interests

The authors declare that they have no competing interests.

## Authors’ contributions

PH, PF, MN, CM, FL, JT, DF, PD, ALL, CB, JBN, AVG, ZP, HK, EW, VN, MS, JAH, TBH, CH,

MK, JS and BHH contributed to the collection/acquisition and logistics of sampling.

MCW, PAM, JAG, WL, YL and EL contributed to the experimental procedures and/or molecular characterization of samples.

MCW, SCE, IRG, SH, NH, AMBP, CF, MAE, PMS, AMA, MK, IG, VN, MS, BHH, JPM and TMB contributed to the data analysis procedures and/or interpretation of results.

MCW, SCE, JMP and TMB contributed to the experimental design of this study. MCW, JMP and TMB wrote the manuscript with feedback from all co-authors.

## Acknowledgments

We are grateful to the Bonobo Conservation Initiative, Vie Sauvage and Institut Congolais pour la Conservation de la Nature, especially Sally Coxe and Albert Lotana Lokasola, for their continuous support of the work of MS. We thank the Ministry of Research of the Democratic Republic of the Congo for permitting the study, and the people of the villages of Bolamba, Yete, Yomboli and Yasalakose for granting access to their forest. We would also like to thank the trackers and scientific assistants of the Manzano forest, and local and national authorities in DRC and the Congolese NGO Mbou-Mon-Tour.

This work was supported by the Spanish funding agency [PID2021-127773NB-I00 funded by MICIU/AEI/10.13039/501100011033 / FEDER/UE; PRE2022-101349 funded by MICIU/AEI /10.13039/501100011033]; grants by the National Institute of Health (R01 AI 050529, R37 AI 150590) to B.H.H., the Bonobo ECO NGO; the Vienna Science and Technology Fund (WWTF) [10.47379/VRG20001] to M.K; the Austrian Science Fund (FWF) [10.55776/ESP546] to S.H.; the computational results of this work have been achieved using supercomputer resources provided by the Life Science Compute Cluster (LiSC) of the University of Vienna; TMB is supported by funding from the European Research Council (ERC) under the European Union’s Horizon 2020 research and innovation programme (grant agreement No. 864203), Grant PID2021-126004NB-100 funded by MICIU/AEI/ 10.13039/501100011033 and ERDF/EU(MICIIN/FEDER, UE) and Revive&Restore Foundation.

## Supplemental Information

### Supplemental Figures

**Fig. S1:**
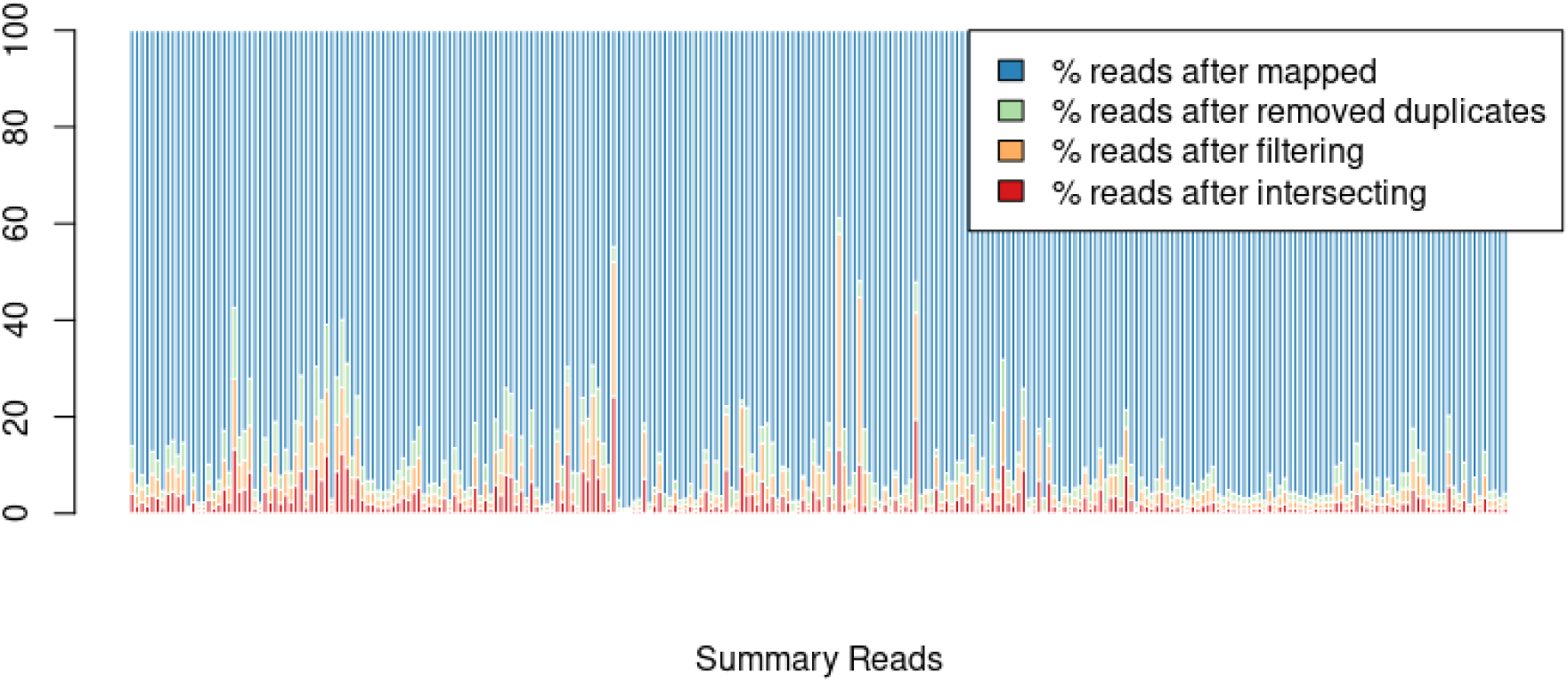
Percentage of reads between each step in our pipeline: blue is the final amount left after mapping, removing duplicates, filtering and intersecting. Image created with R (Posit Team 2022; R Core Team software Version 4.4.x. 2025).

**Fig. S2:**
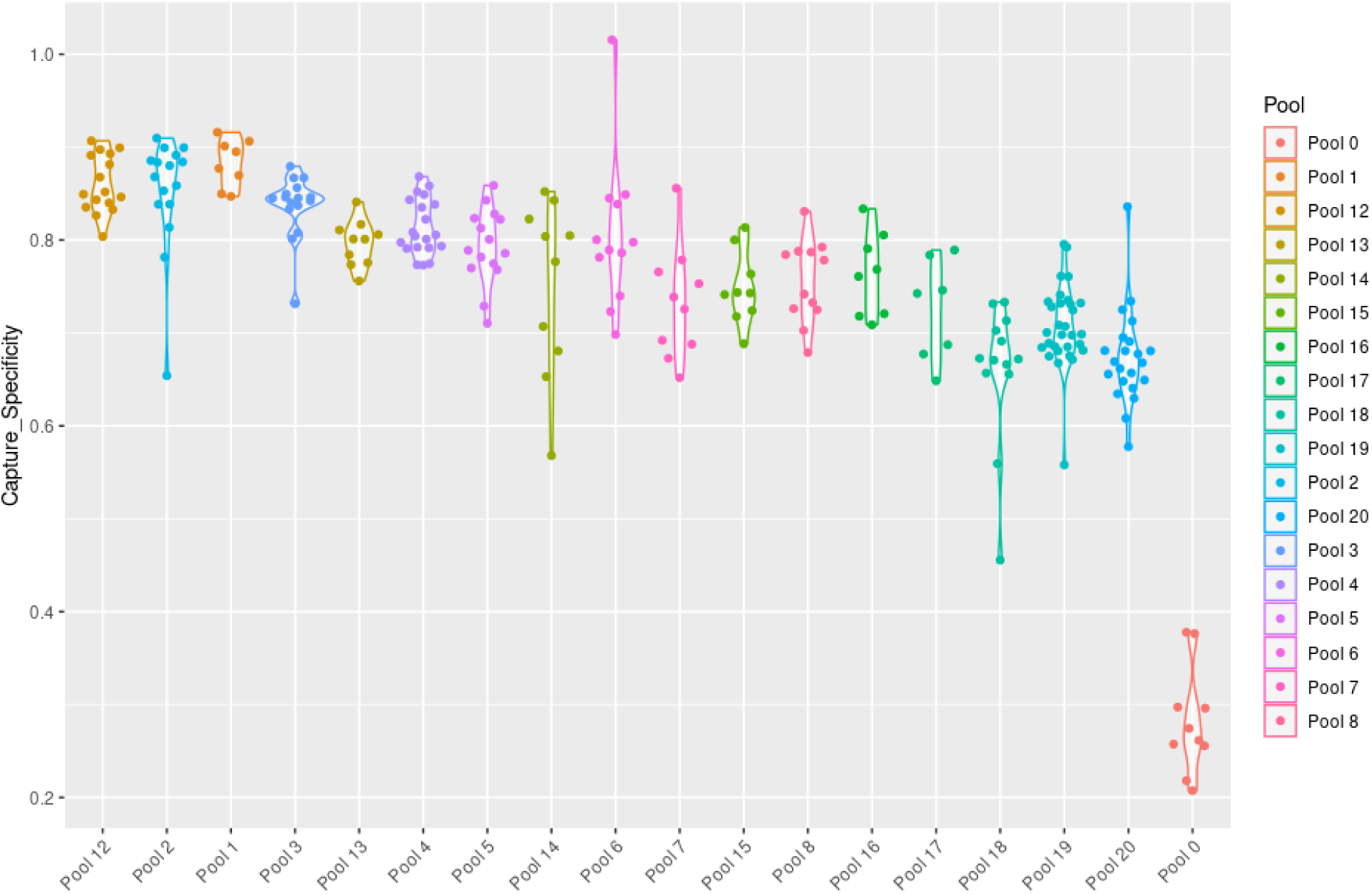
Capture specificity of each pool, calculated by dividing reliable reads on-target by reliable reads. Image created with R (Posit Team 2022; R Core Team software Version 4.4.x. 2025).

**Fig. S3:**
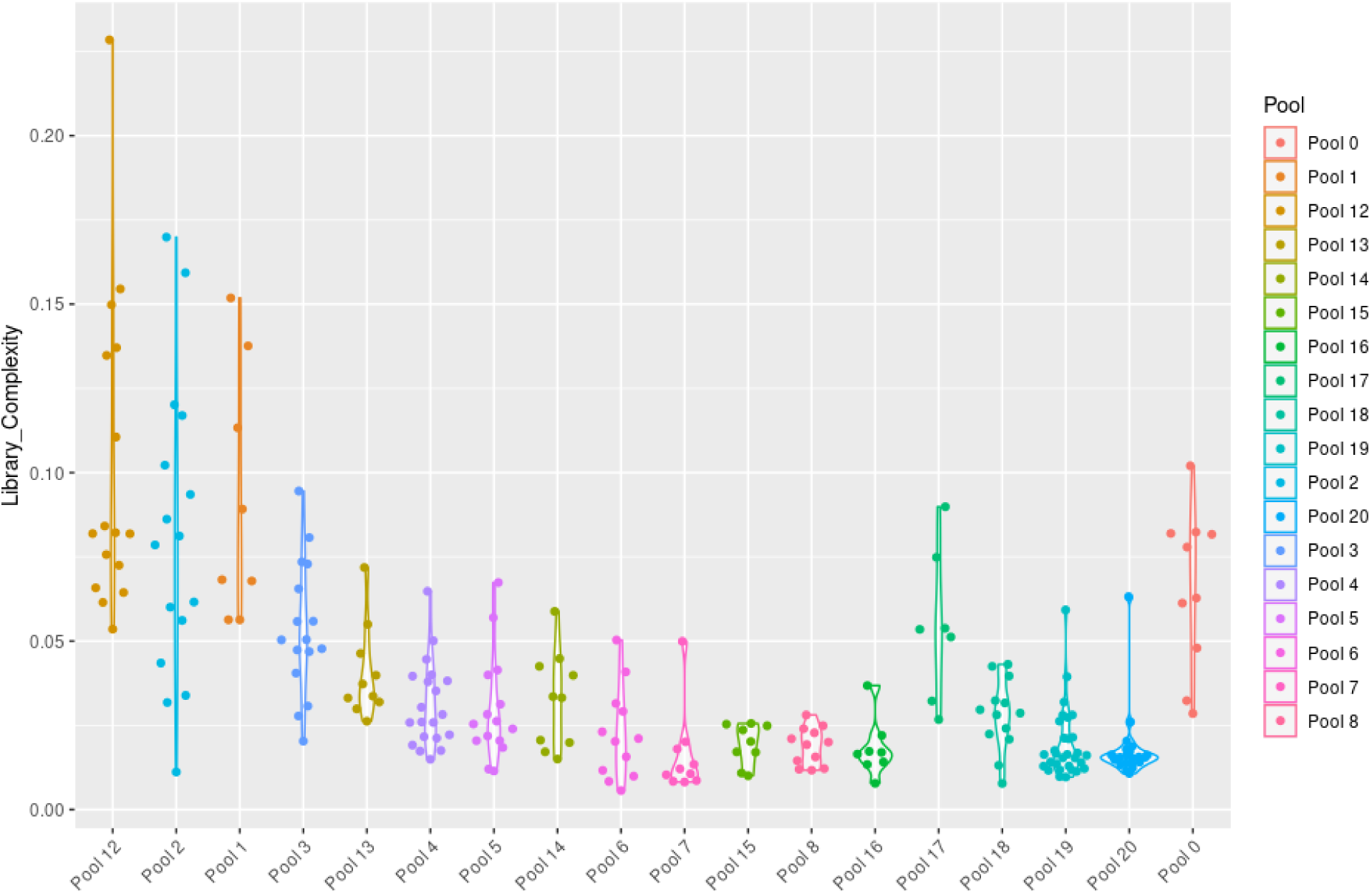
Library complexity of each pool, calculated by dividing reliable reads by mapped reads. Image created with R (Posit Team 2022; R Core Team software Version 4.4.x. 2025).

**Fig. S4:**
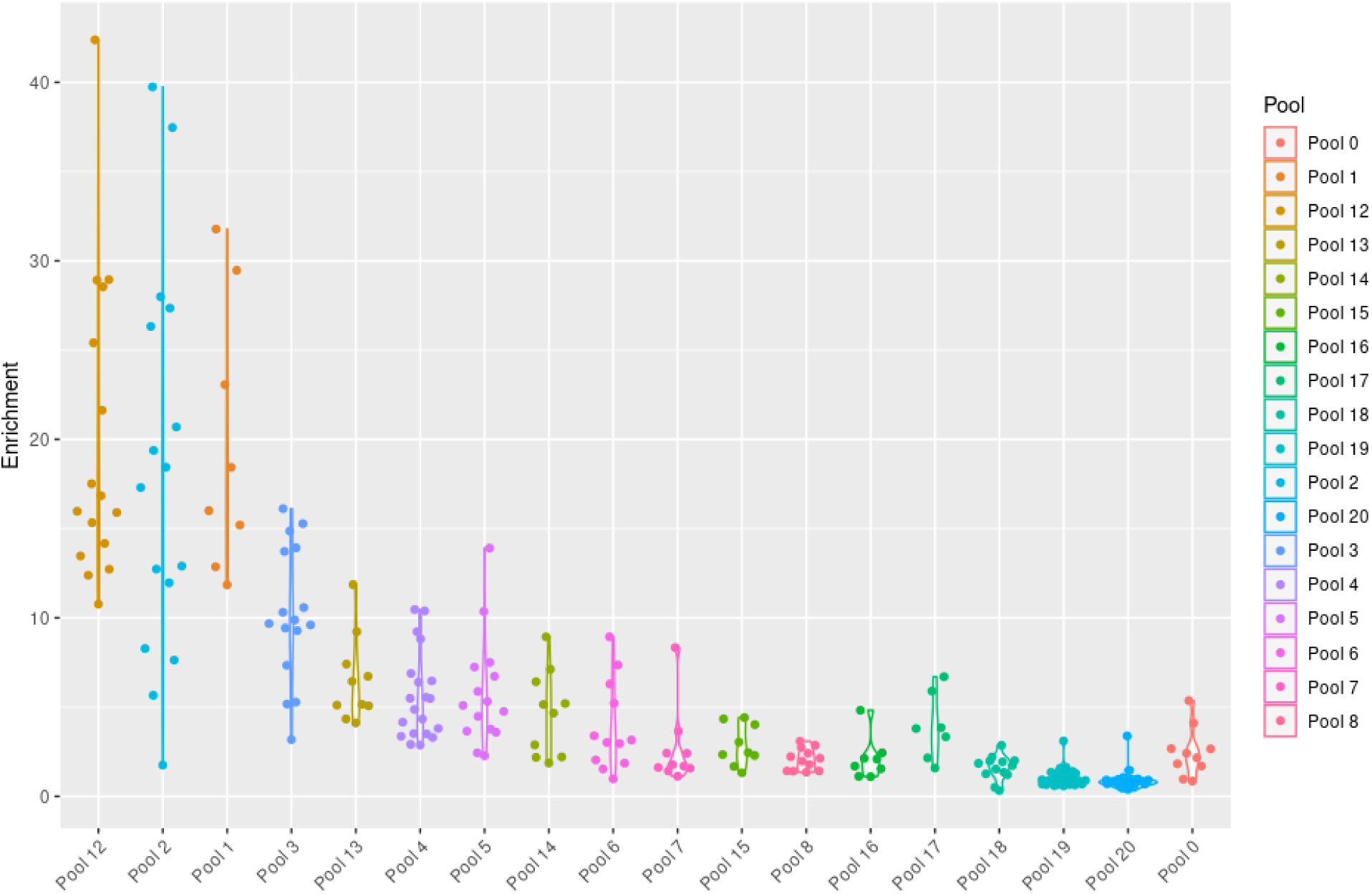
Enrichment performance of each pool, calculated by the following: (reliable reads on-target/by mapped reads)/(chr21 reads/genome size). Image created with R (Posit Team 2022; R Core Team software Version 4.4.x. 2025).

**Fig. S5:**
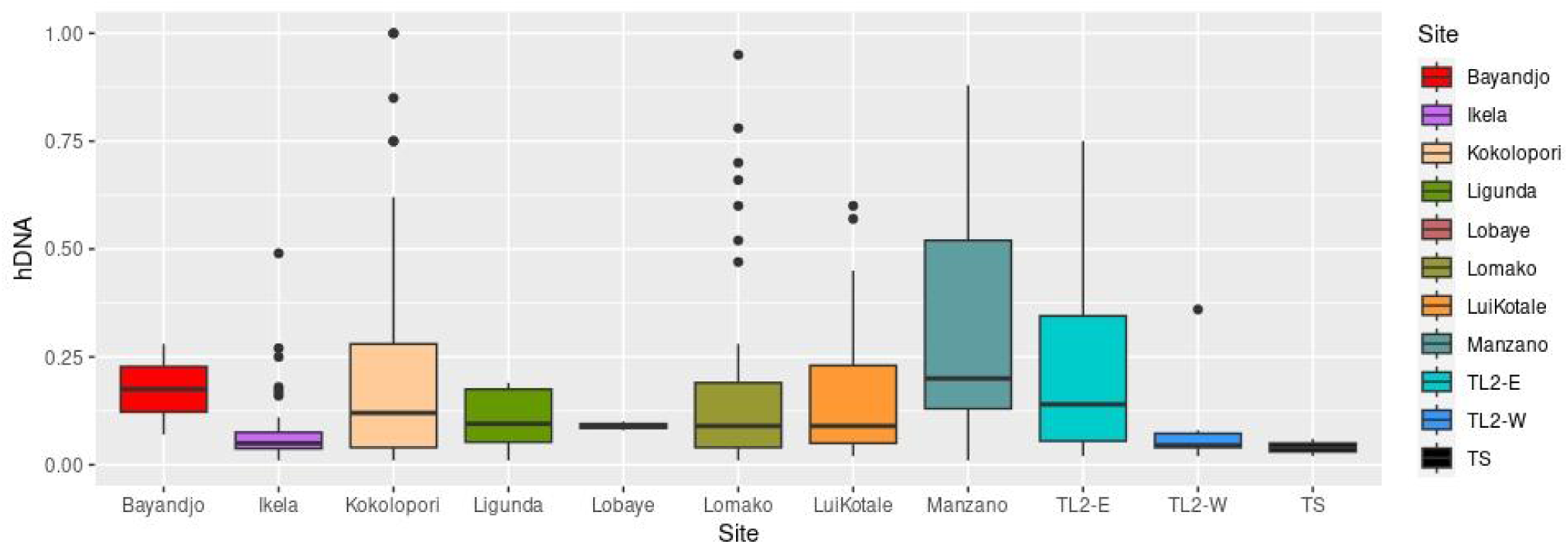
Boxplot of percentage host DNA in each site prior to sequencing. Image created with R (Posit Team 2022; R Core Team software Version 4.4.x. 2025).

**Fig. S6:**
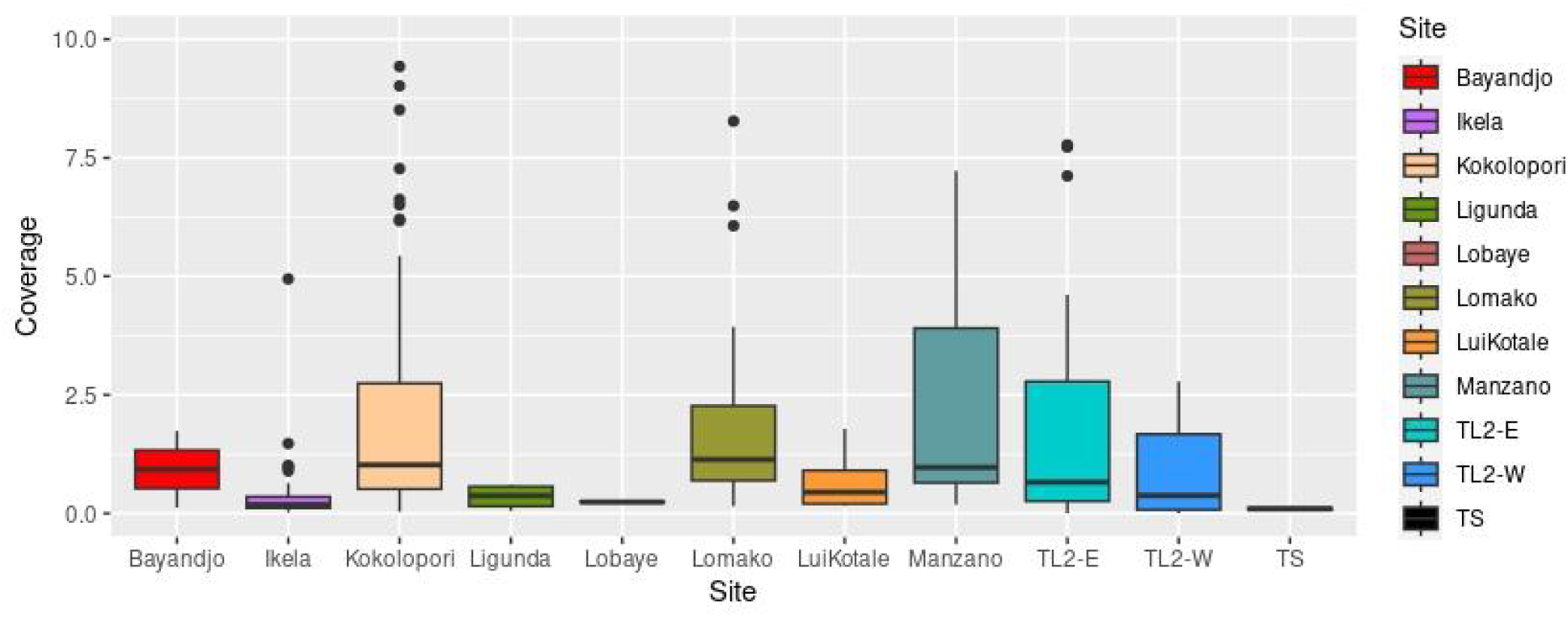
Boxplot of depth of coverage in each site after filtering at ≥0.1x and 1% human contamination. Image created with R (Posit Team 2022; R Core Team software Version 4.4.x. 2025).

**Fig. S7:**
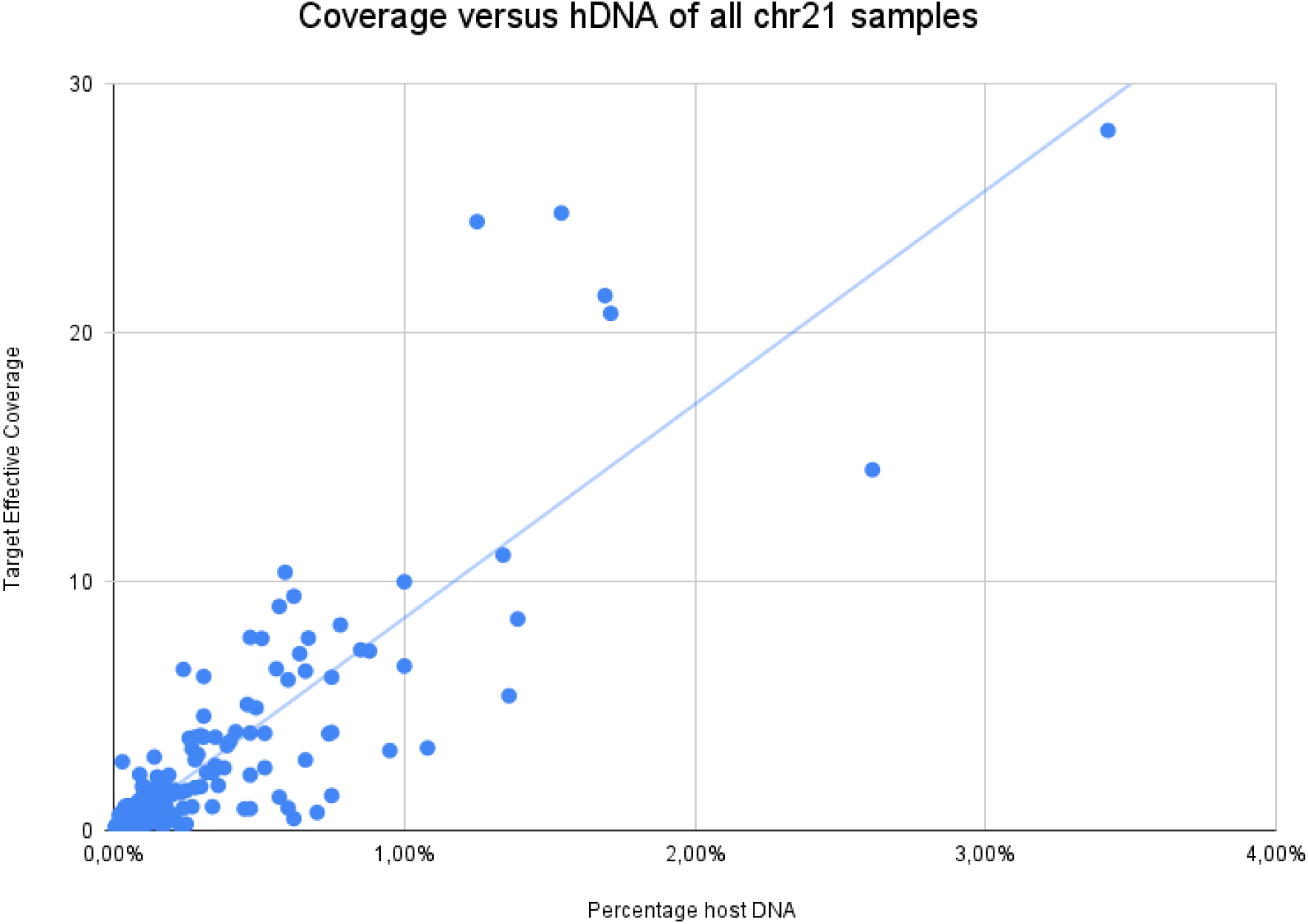
Scatterplot of depth of coverage versus host DNA per sample. Image created with Microsoft Excel (Microsoft Corporation 2018).

**Fig. S8:**
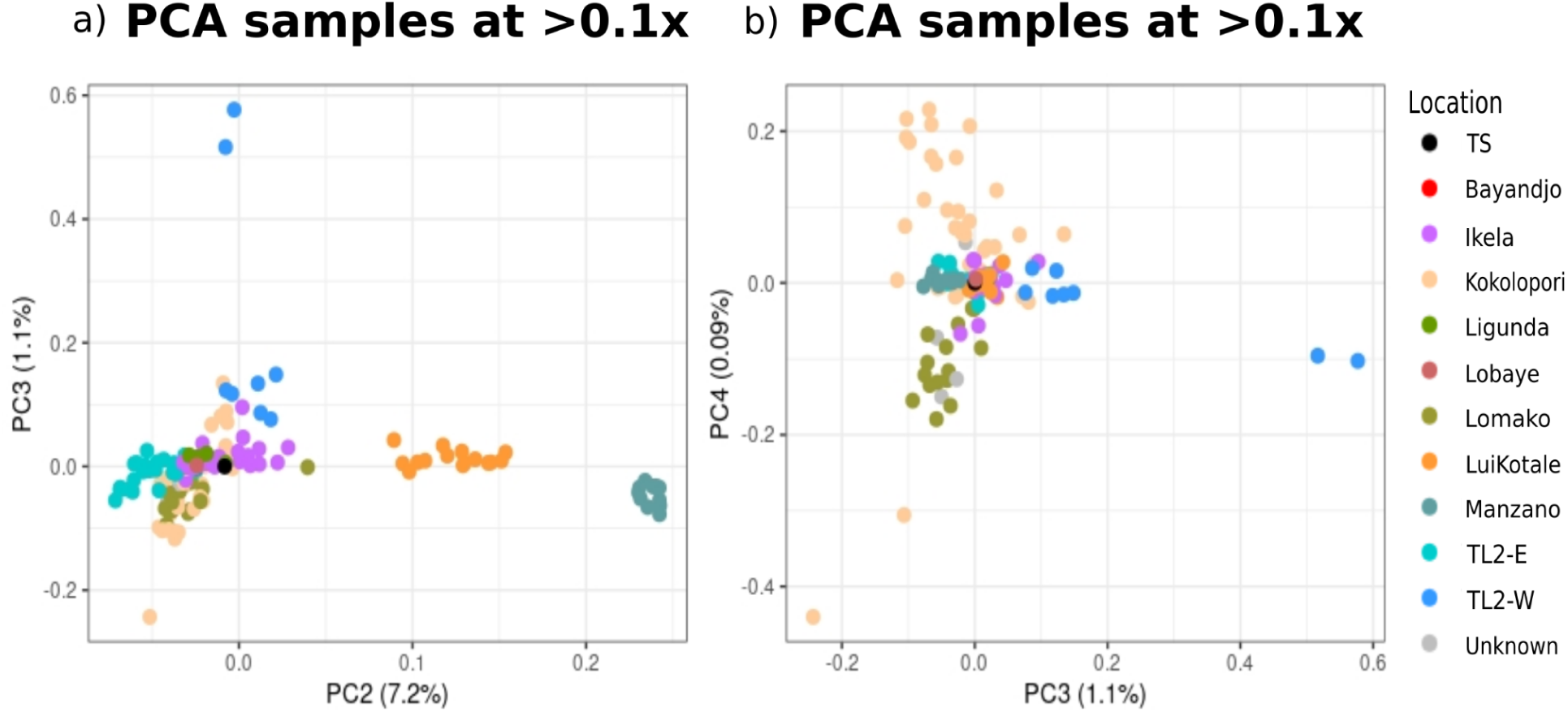
Principal Component Analysis: a) components 2 and 3 using PCAngsd of all samples filtered at ≥0.1x & 1% human contamination; b) components 3 and 4 using PCAngsd of all samples filtered at ≥0.1x & 1% human contamination (Meisner & Albrechtsen 2018). Some samples in grey were mislabelled but have been identified as Lomako and Kokolopori. Image created with R (Posit Team 2022; R Core Team software Version 4.4.x. 2025).

**Fig. S9:**
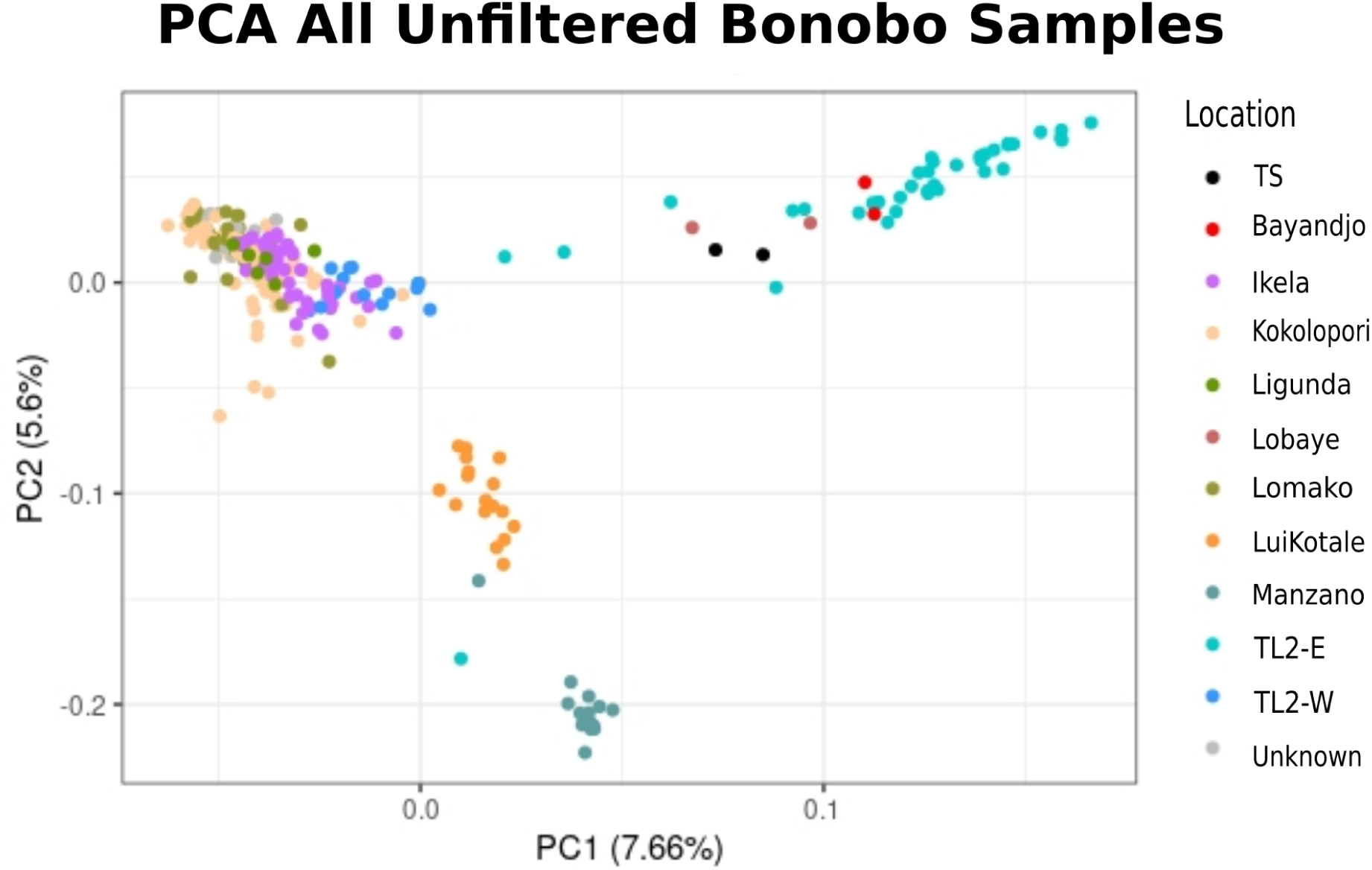
Principal Component Analysis using PCAngsd (Meisner & Albrechtsen 2018) of all samples unfiltered. Some samples in grey were mislabelled but have been identified as Lomako and Kokolopori. Image created with R (Posit Team 2022; R Core Team software Version 4.4.x. 2025).

**Fig. S10:**
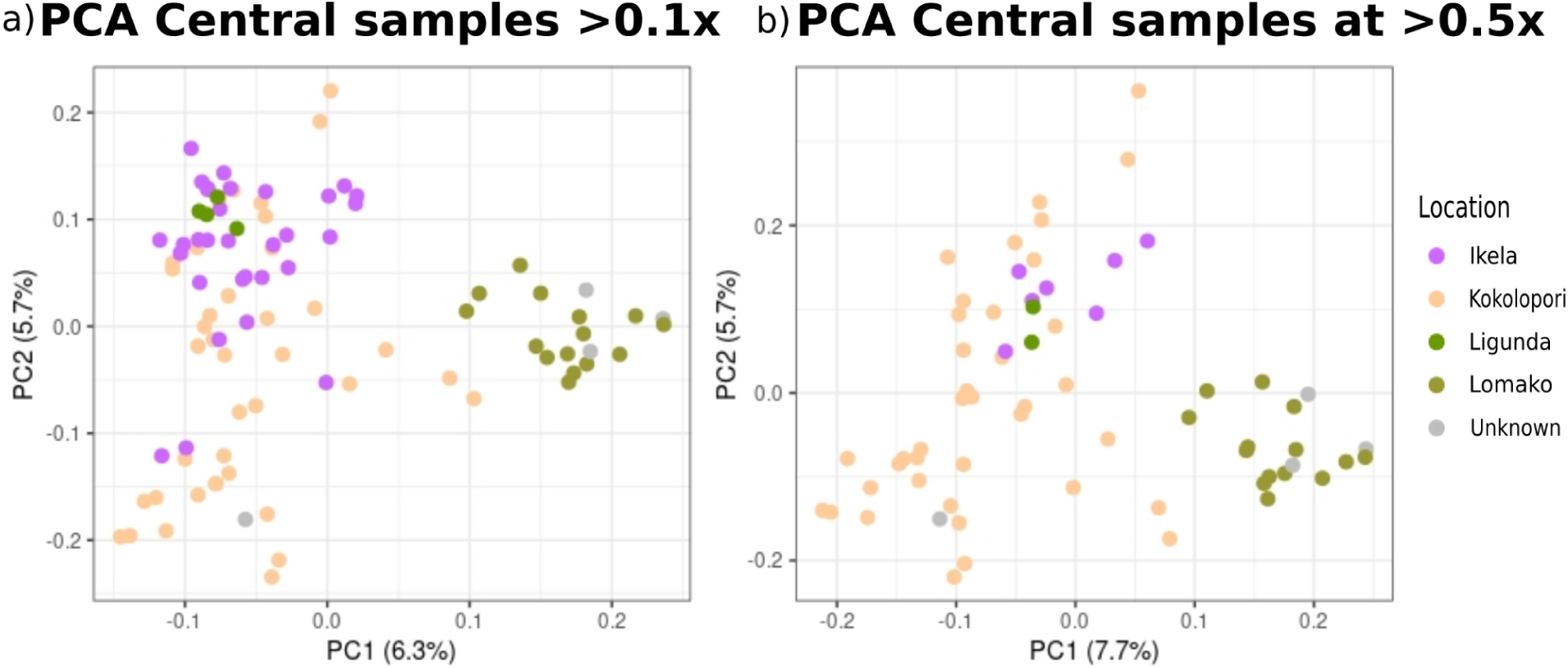
Principal Component Analysis using PCAngsd (Meisner & Albrechtsen 2018): a) North-Central bonobo samples without TL2-W and filtered at ≥0.1x & 1% human contamination; b) North-Central bonobo samples without TL2-W and filtered at ≥0.5x & 1% human contamination. Central samples here are from Ikela, Kokolopori, Ligunda, Lomako and samples that were mislabelled but have been identified as Lomako and Kokolopori. Image created with R (Posit Team 2022; R Core Team software Version 4.4.x. 2025).

**Fig. S11:**
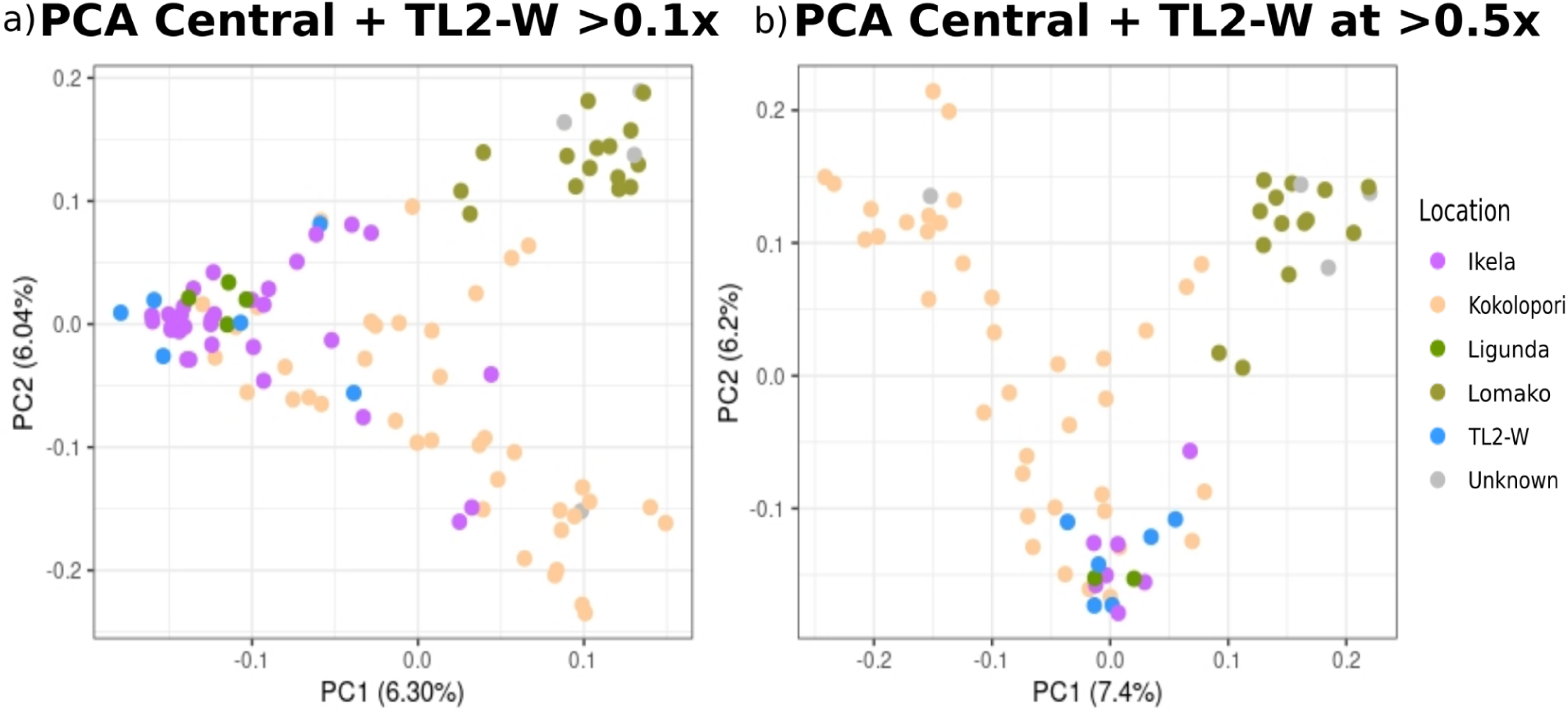
Principal Component Analysis using PCAngsd (Meisner & Albrechtsen 2018): a) North-Central bonobo samples and TL2-W, and filtered at ≥0.1x & 1% human contamination; b) North-Central bonobo samples and TL2-W, and filtered at ≥0.5x & 1% human contamination. Samples are from TL2-W, Ikela, Kokolopori, Ligunda, Lomako and samples that were mislabelled but have been identified as Lomako and Kokolopori. Image created with R (Posit Team 2022; R Core Team software Version 4.4.x. 2025).

**Fig. S12:**
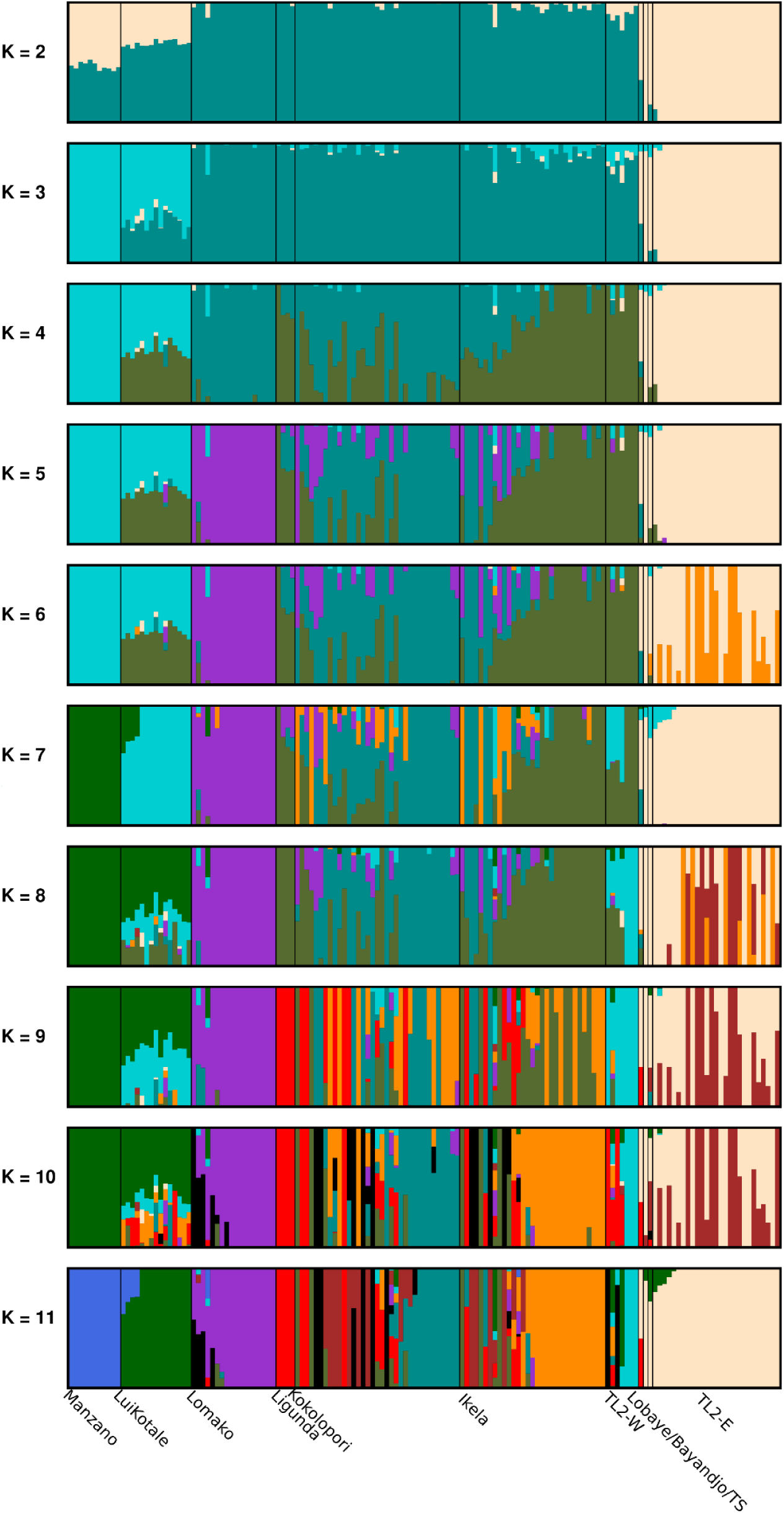
bar-plot of admixture values per individual grouped per site at ≥ 0.1x depth of coverage of first run from K2 to K11 without whole genome samples. Image created with pong (Behr et al. 2016).

**Fig. S13:**
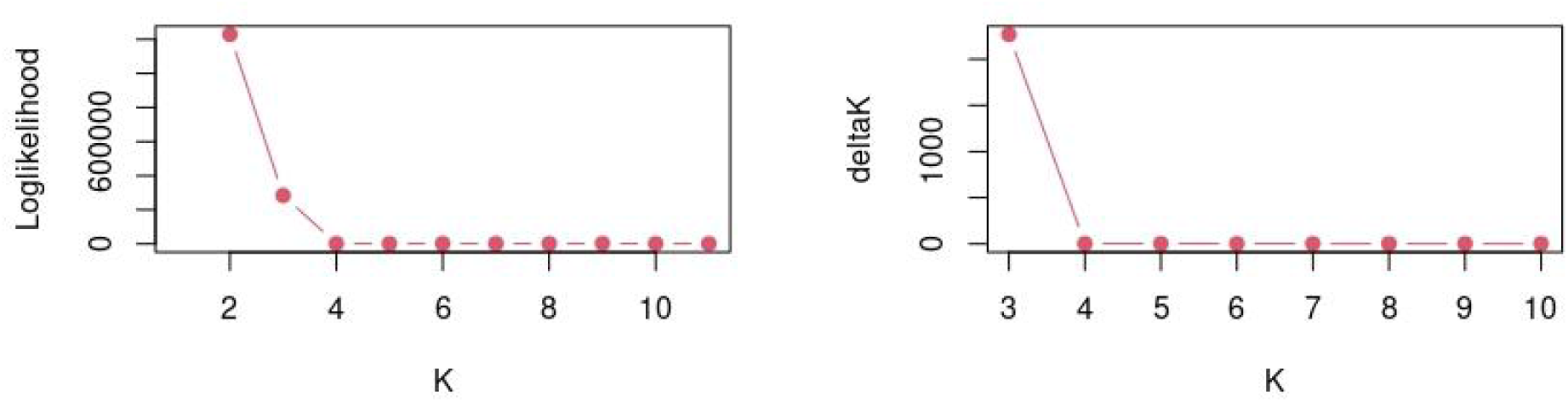
NgsAdmix plot for inferred K: left is the inferred K calculated from the normalized mean of likelihoods from all Ks; right is the inferred K calculated from the normalized mean of differences from likelihoods of Ks. Image created with R (Posit Team 2022; R Core Team software Version 4.4.x. 2025).

**Fig. S14:**
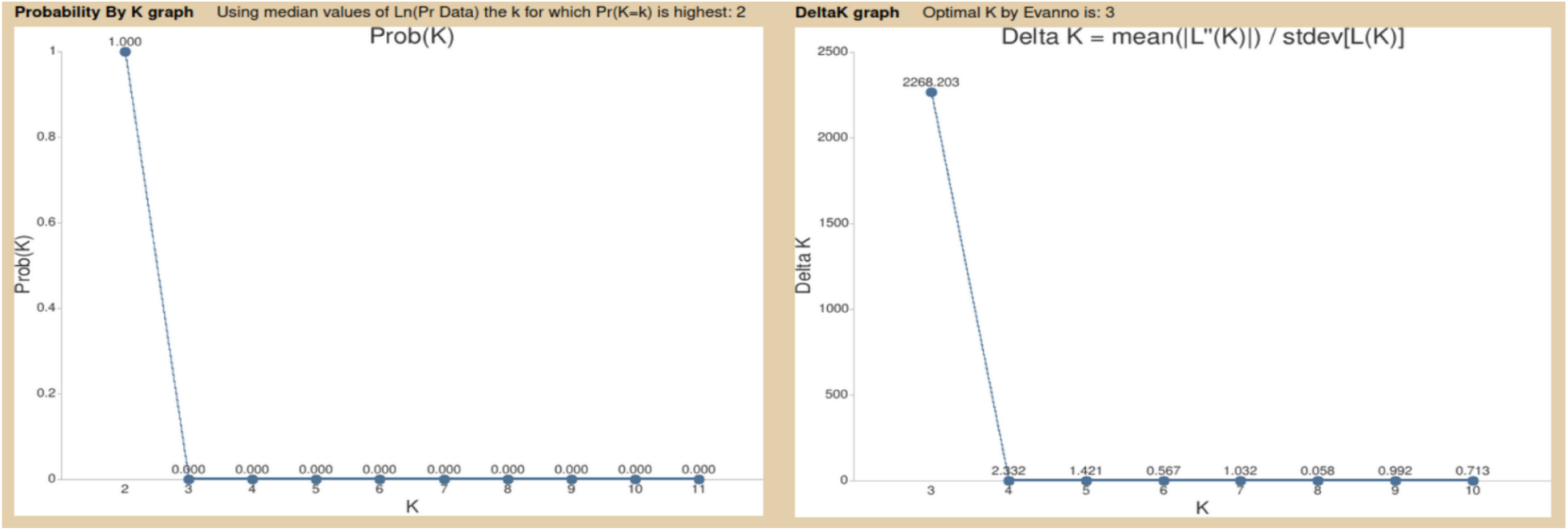
Clumpak plot for inferred K: left is the inferred K calculated from the normalized mean from all Ks; right is the inferred K calculated from the normalized mean of differences from likelihood Ks. Output image created by Clumpak (https://tau.evolseq.net/clumpak/index.html).

**Fig. S15:**
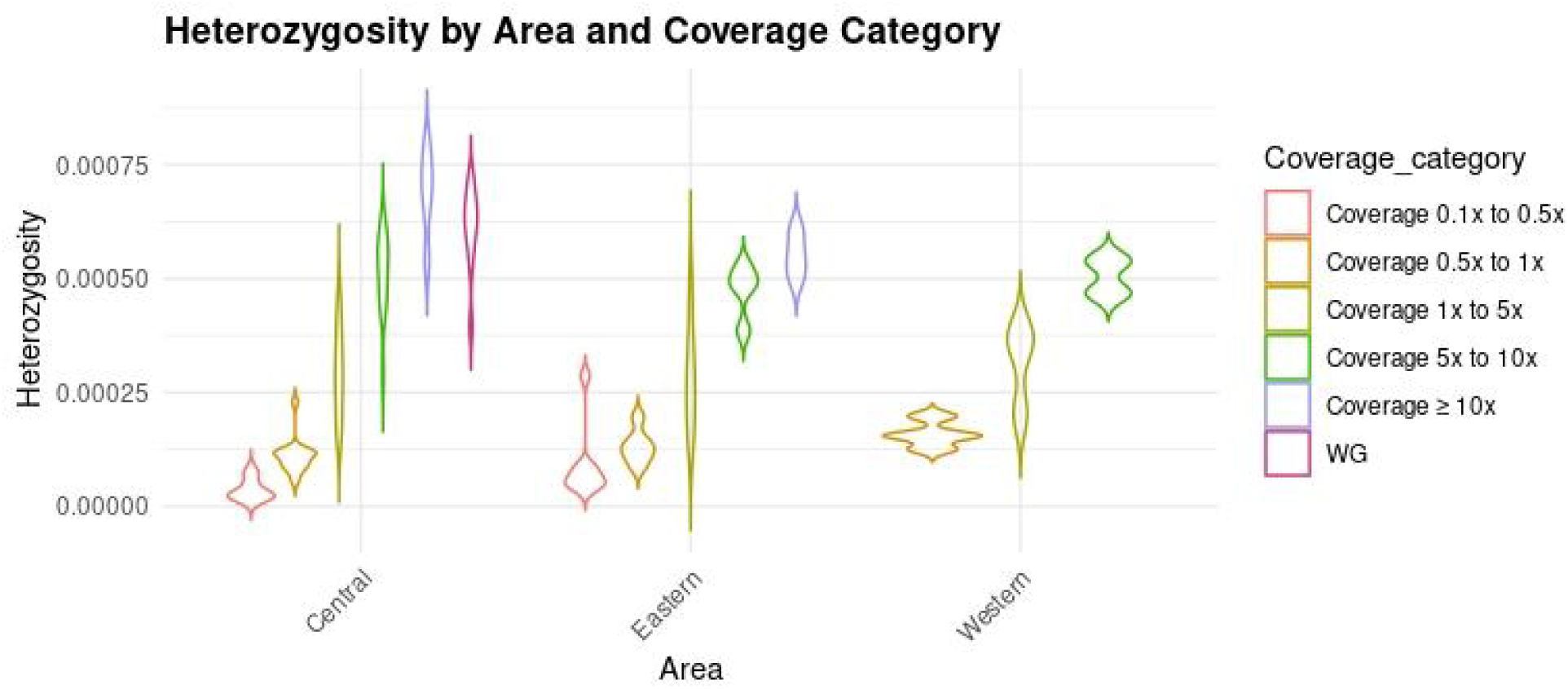
Violin plot of heterozygosity levels per area at different depths of coverages with whole genome bonobo samples (WG) from the Prado et al. 2013 paper. Image created with R (Posit Team 2022; R Core Team software Version 4.4.x. 2025).

**Fig. S16:**
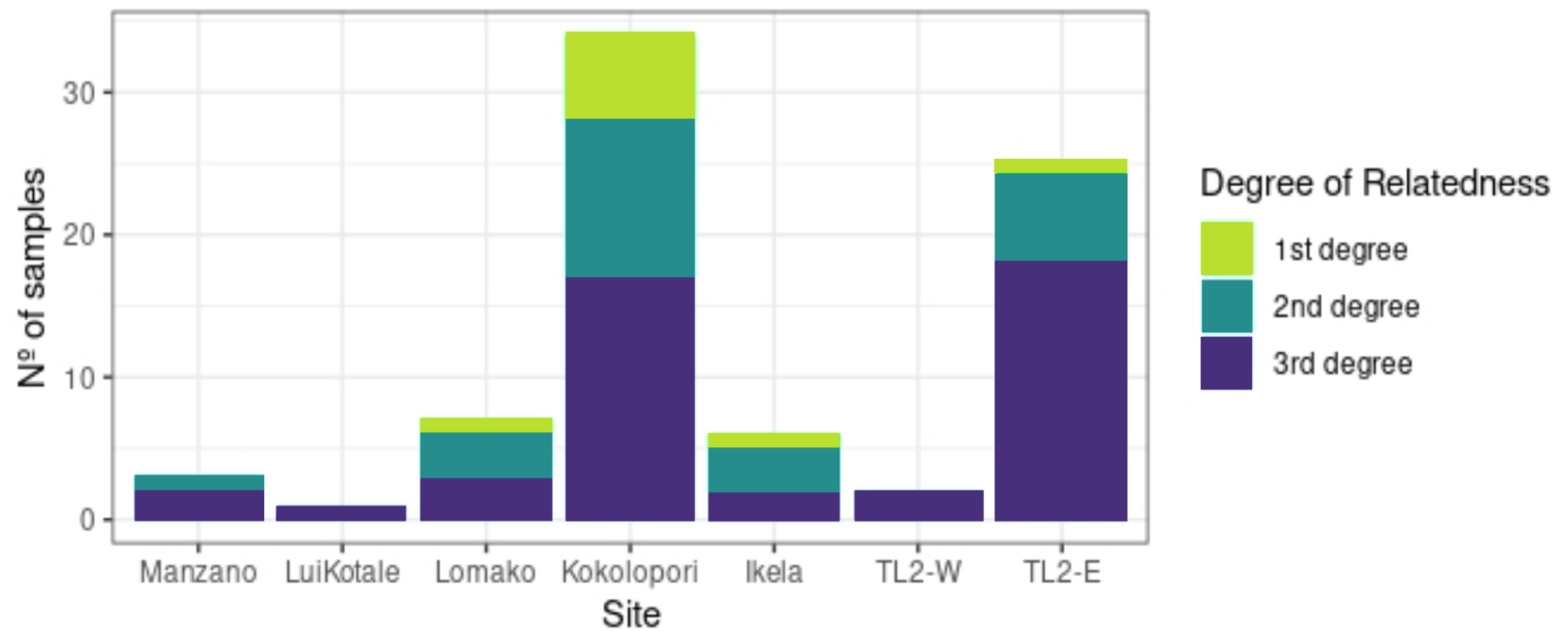
Barplot of total number of related samples for each site at 0.3x depth of coverage or more. Sample size of first degree for each site: Lomako (n=1); Kokolopori (n=6); Ikela (n=1); TL2-E (n=1). Image created with R (Posit Team 2022; R Core Team software Version 4.4.x. 2025).

**Fig. S17:**
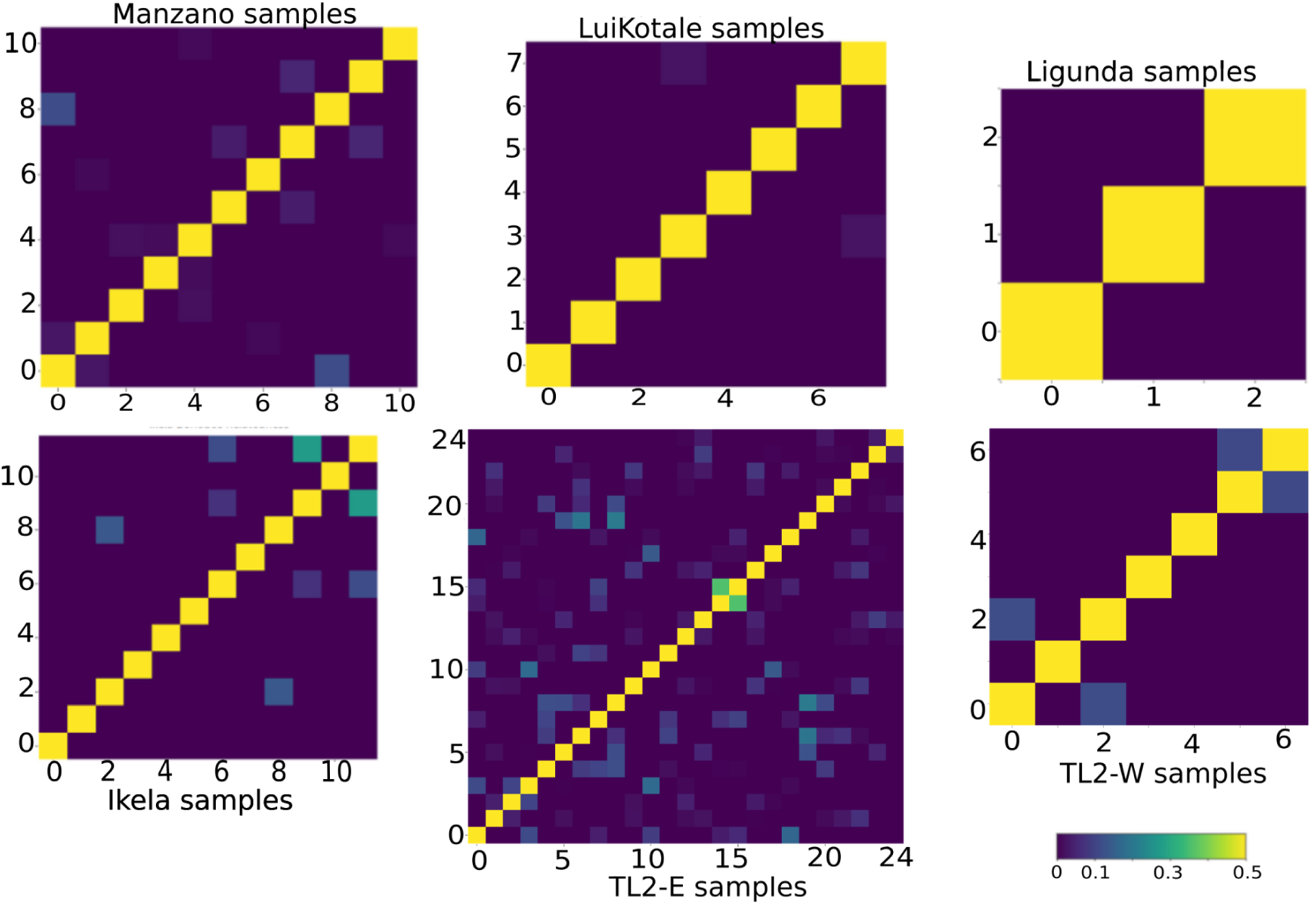
Relatedness-site matrices made using NgsRelate v.2 (Korneliussen et al. 2015) of all bonobo samples filtered, QC passed and with depth of coverage above 0.3x. Axis represents the number of samples and the gradient is level of relatedness. Closer to yellow is a higher level whilst yellow represents the same individual. Images were created with R (Posit Team 2022; R Core Team software Version 4.4.x. 2025).

**Fig. S18:**
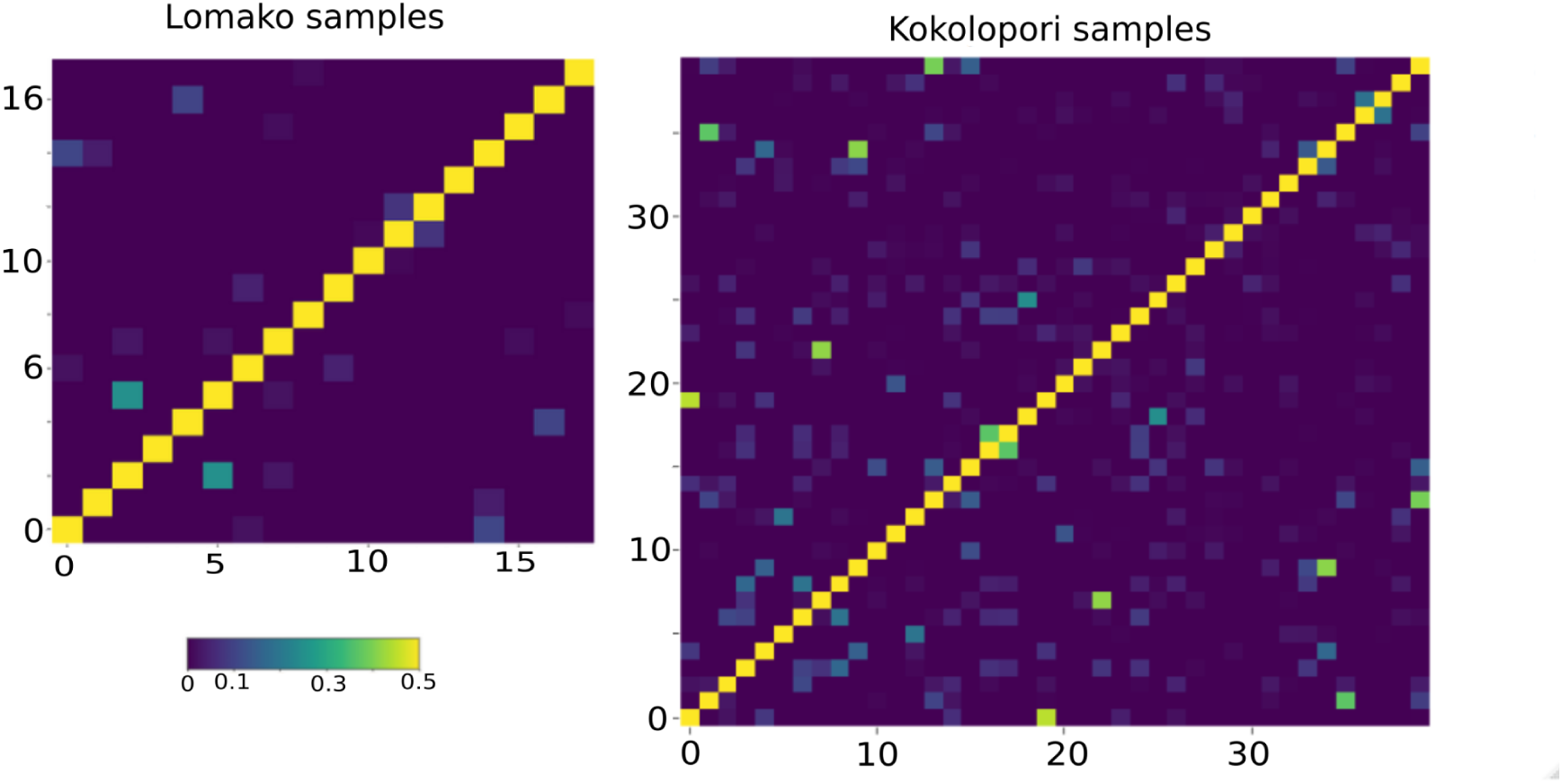
Site matrices of LuiKotale and Kokolopori samples made using NgsRelate v.2 (Korneliussen et al. 2015) of all bonobo samples filtered, QC passed and with depth of coverage above 0.3x. Axis represents the number of samples and the gradient is level of relatedness. Closer to yellow is a higher level whilst yellow represents the same individual. Images were created with R (Posit Team 2022; R Core Team software Version 4.4.x. 2025).

**Fig. S19:**
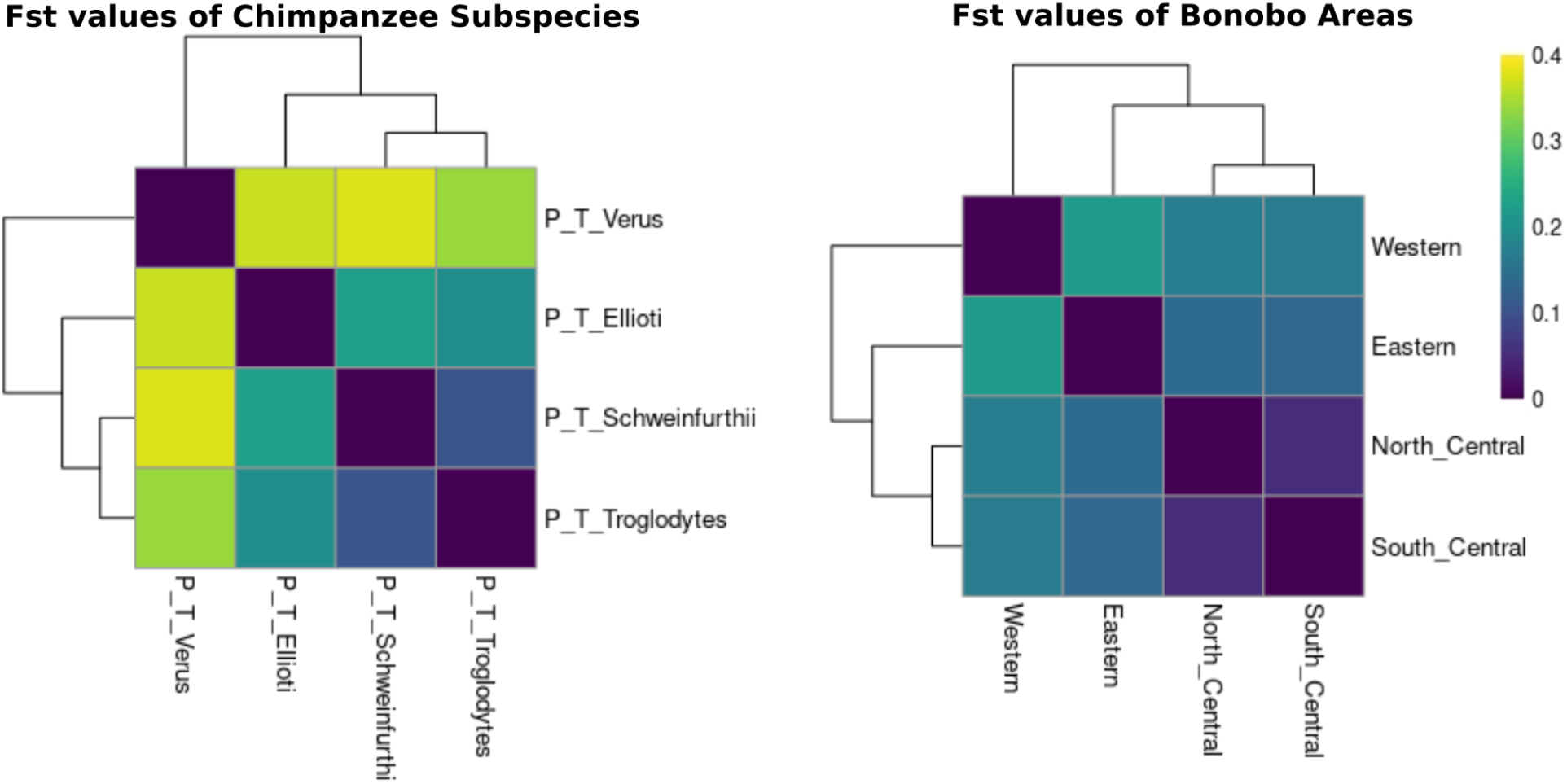
Heat-maps of *F_ST_*indexes: left are values for chimpanzee subspecies; right are values for bonobo areas. Barriers such as rivers were not taken into account as we use large groups of sites and subspecies. Bonobo areas are here defined as: Western = Manzano samples, Eastern = TL2-E samples, North Central = Lomako and Kokolopori samples, South Central = Ikela and TL2-W samples. Darker colouring indicates lower genetic differentiation whilst the lighter yellow colouring indicates higher genetic variation. Image created with R (Posit Team 2022; R Core Team software Version 4.4.x. 2025).

**Fig. S20:**
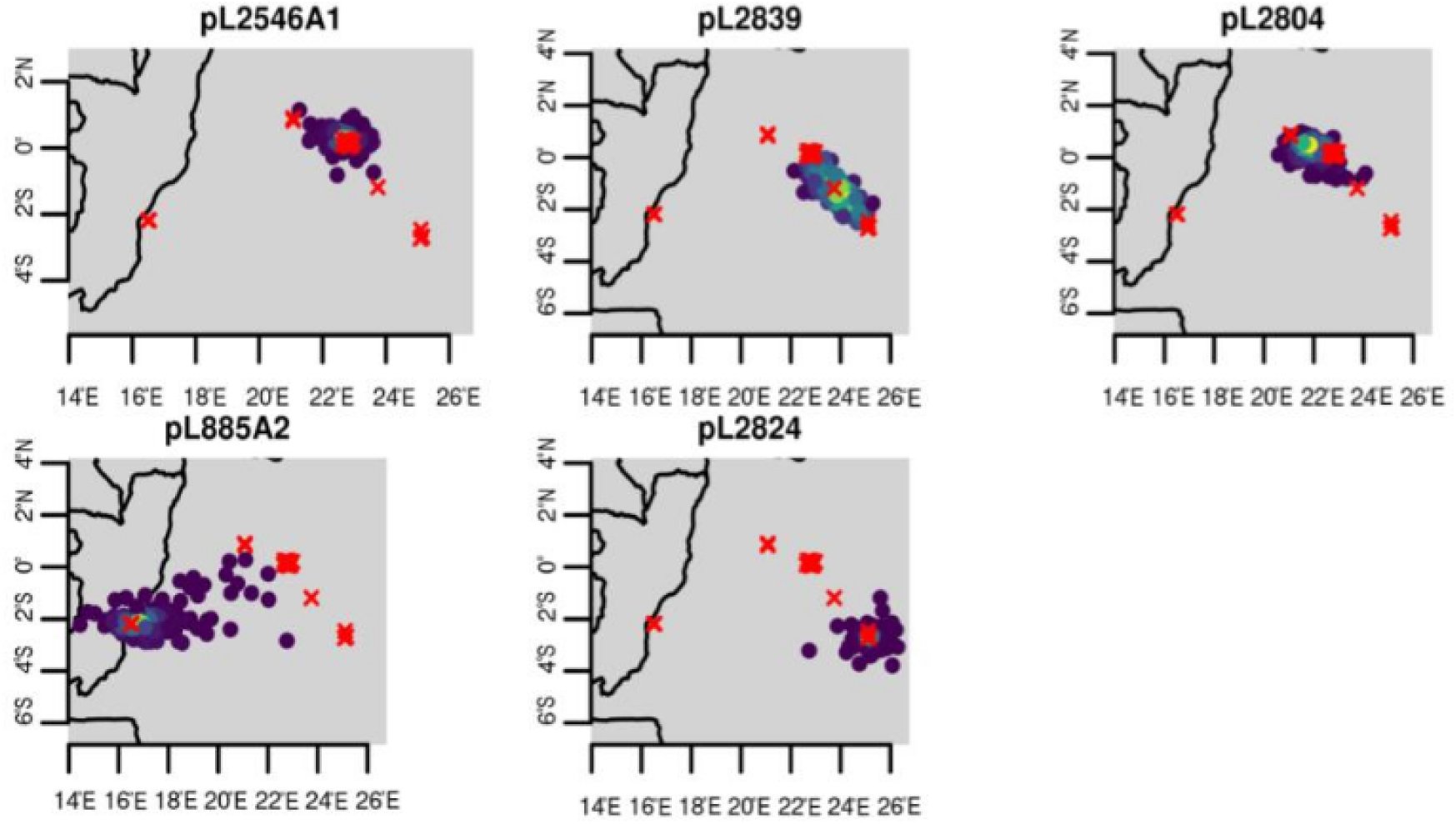
Output test plots of Locator data with real coordinates used for reference (red crosses) and predicted location (circles, with colour grading according to number of coinciding predictions - lighter ones being the most probable and darker least). Samples’ real origin is: pL2546A1 - Kokolopori; pL2839 - TL2-W; pL2804 - Lomako; pL885A2 - Manzano; pL2824 - TL2-E. Image edited with R (Posit Team 2022; R Core Team software Version 4.4.x. 2025).

**Fig. S21:**
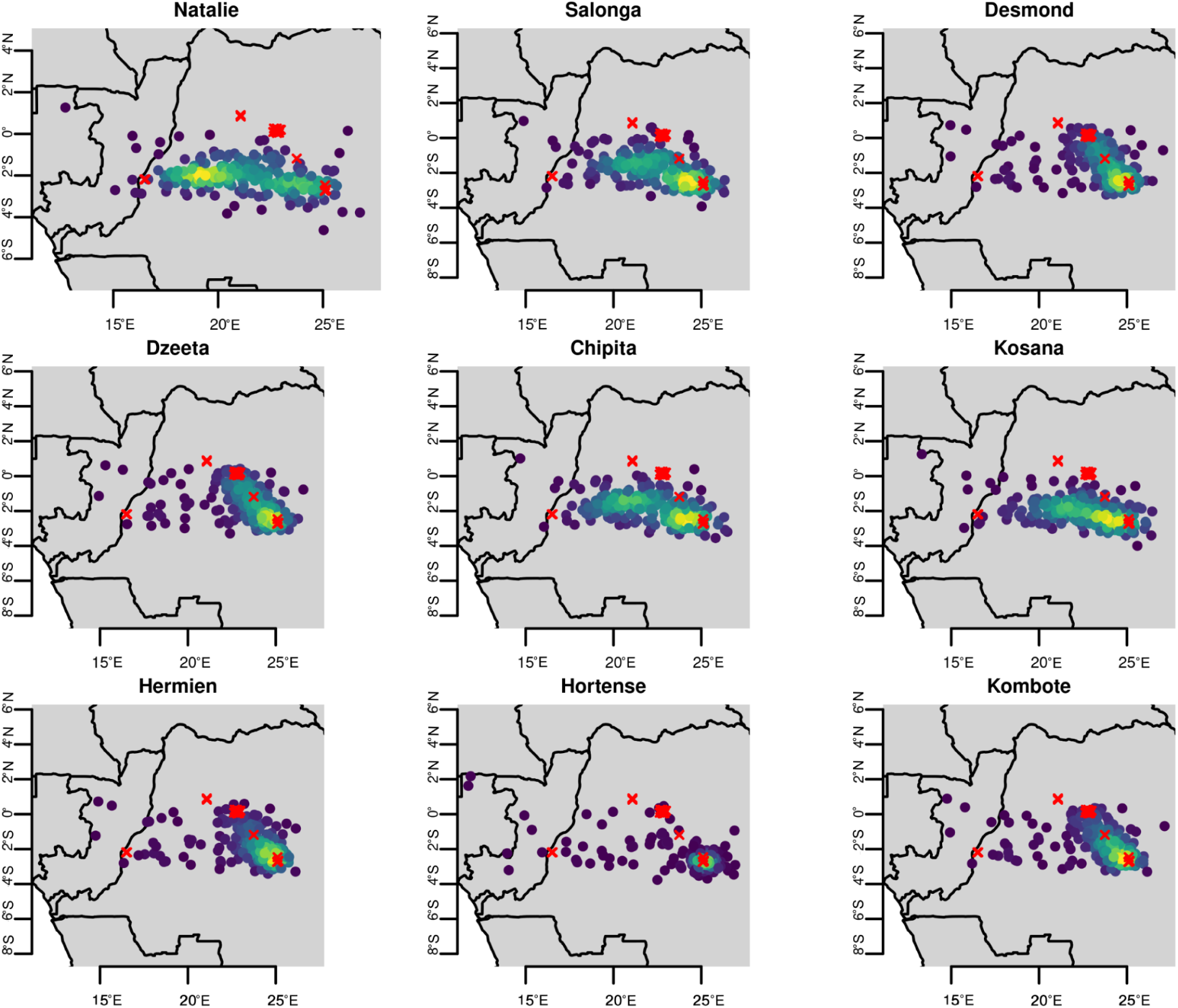
Output plots of Locator data with real data coordinates used for reference (red crosses) and predicted location (circles, with colour grading according to number of coinciding predictions - lighter ones being the most probable and darker the least).

**Fig. S22:**
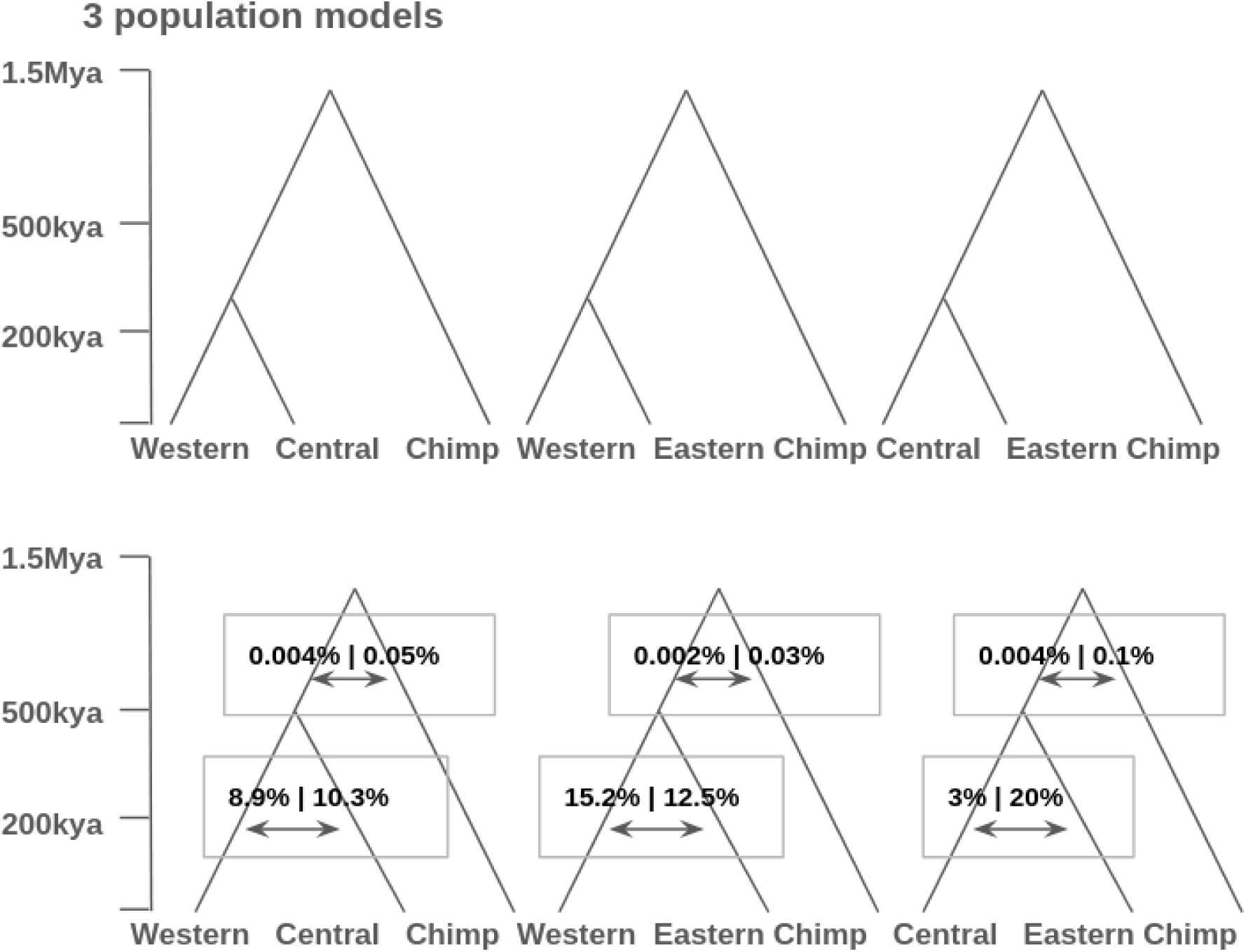
A schematic representation of the G-PhoCS analyses. Each analysis included two bonobo populations and a Western chimpanzee lineage as an outgroup. We considered three models without migration (top) and three models with migration (bottom). Models with migration included four directional migration bands, as depicted in the figure. The scaled inferred divergence times are depicted by the height of the corresponding split in the tree (see y-axis). Inferred migration rates in % are written on the arrows, e.g. 0.004% from bonobos to chimpanzees and 8.9% from western to central bonobos on the WeCeChimp model on the bottom left. See Table S4 for the estimated values of these parameters.

## Notes

### Competing Interest Statement

The authors have declared no competing interest.

